# MCUR1–CCDC90B complex is a conserved mitochondrial scaffold regulating metabolic homeostasis

**DOI:** 10.1101/2025.10.14.682030

**Authors:** J Verissimo Ferreira, J Wettmarshausen, V Goh, K Lagerborg, J Watrous, MS Feng, H Delgado de la Herran, S Walia, Y Cheng, A Leimpek, M Gorza, S Braun, R Nilsson, M Jain, M Mann, D Mokranjac, M Murgia, TB Haack, M Deschauer, H Prokisch, FV Filipp, F Perocchi

**Affiliations:** Institute for Diabetes and Obesity, Helmholtz Diabetes Center, Helmholtz Munich, Munich, Germany; Departments of Medicine and Pharmacology, UC San Diego School of Medicine, La Jolla, CA, USA; Institute for Genetics, Justus-Liebig-University Giessen, Giessen, Germany; Department of Medicine, Karolinska Institute, Karolinska University Hospital, Stockholm, Sweden; Department of Proteomics and Signal Transduction, Max Planck Institute of Biochemistry, Martinsried, Germany; Faculty of Health Sciences, Novo Nordisk Foundation Center for Protein Research, University of Copenhagen, Copenhagen, Denmark; LMU Munich, Biozentrum-Cell Biology, Planegg-Martinsried, Germany; BioMedical Center (BMC), Division of Physiological Chemistry, Faculty of Medicine, LMU Munich, Planegg-Martinsried, Germany; Department of Biomedical Sciences, University of Padova, Padua, Italy; Institute of Medical Genetics and Applied Genomics, University of Tübingen, Tübingen, Germany; Centre for Rare Diseases, University of Tuebingen, Tübingen, Germany; Department of Neurology, Klinikum rechts der Isar, School of Medicine and Health, Technical University of Munich, Munich, Germany; Institute of Neurogenomics, Computational Health Center, Helmholtz Munich, Munich, Germany; School of Medicine, Institute of Human Genetics, Technical University of Munich, Munich, Germany; School of Life Sciences Weihenstephan, Technical University of Munich, Freising, Germany; Institute for Advanced Study, Technical University of Munich, Munich, Germany; Metaflux, San Diego, CA, USA; German Center for Diabetes Research, Munich, Germany; Institute of Neuronal Cell Biology, Technical University of Munich, Munich, Germany; Munich Cluster for Systems Neurology, Munich, Germany

**Keywords:** MCUR1, CCDC90B, mitochondria, mitochondrial metabolism, amino acid metabolism

## Abstract

The mitochondrial calcium uniporter regulator 1 (MCUR1) is an evolutionarily conserved protein of the inner mitochondrial membrane^1^, yet its physiological role has remained elusive. Although initially proposed to function as a subunit of the mitochondrial calcium uniporter complex (MCUC)^2–4^, emerging evidence suggests that MCUR1 has a broader functional spectrum^5–7^. Here, we identify a biallelic loss-of-function *MCUR1* variant (c.802C>T; p.R268X) in a patient with a progressive neurological phenotype. This mutation leads to loss of MCUR1 protein and exerts a dominant-negative effect on its paralog, CCDC90B. We show that MCUR1 and CCDC90B form a hetero-oligomeric complex whose stability depends on MCUR1. Deletion of *MCUR1* and *CCDC90B* in the fission yeast *Schizosaccharomyces pombe*, which lacks the MCUC, impairs lipid and amino acid metabolism and causes nitrogen source-dependent growth defects that are rescued by expression of human MCUR1. Patient serum metabolomics confirms an imbalance in the amino acid pool, while *MCUR1* deficiency in patient-derived skin fibroblasts upregulates autophagy, perturbs non-essential amino acid metabolism, and limits biosynthetic capacity, resulting in delayed proliferation and migration. These findings redefine the MCUR1–CCDC90B coiled-coil complex as a transmembrane scaffold critical for the integrity of mitochondrial protein complexes and the maintenance of metabolic homeostasis, suggesting a potential link between MCUR1 deficiency and human neurometabolic disease.

## INTRODUCTORY PARAGRAPH

MCUR1 (previously annotated as CCDC90A) and its paralog CCDC90B belong to a functionally heterogeneous family of coiled-coil domain-containing (CCDC) proteins implicated in scaffolding, membrane fusion, and signal transduction^1^. Despite strong evolutionary conservation, the function of MCUR1 and CCDC90B has remained unclear. Loss of MCUR1 in mammalian cell lines and heart mitochondria impairs mitochondrial Ca^2+^ (mt-Ca^2+^) uptake by destabilizing MCUC assembly^2–4^. By contrast, overexpression of human MCUR1 in *Drosophila* cells, which lack MCUR1 homologs, reduces mt-Ca^2+^ buffering capacity and sensitizes mitochondria to Ca^2+^-induced permeability transition, likely through a direct interaction with components of the mitochondrial permeability transition pore^6^. Subsequent studies in human fibroblasts and in the budding yeast *Saccharomyces cerevisiae* have challenged this Ca^2+^-centric model, proposing instead that MCUR1 regulates the assembly of respiratory chain complex (RCC) IV (cytochrome c oxidase)^5^ or influences proline utilization^7^. The reported effects of MCUR1 loss-of-function on mt-Ca^2+^ uptake may therefore reflect perturbations in cellular bioenergetics and mitochondrial membrane potential. Collectively, these observations raise fundamental questions about the primary function of MCUR1 and its potential relevance to human physiology.

Here, we report the first identification of a patient with a loss-of-function variant in *MCUR1* that abolishes the expression of both MCUR1 and CCDC90B. Through integrative systems analyses in patient-derived fibroblasts, HeLa cells, and two evolutionarily distant yeast models (*Saccharomyces cerevisiae* and *Schizosaccharomyces pombe)*, we demonstrate that MCUR1 is dispensable for mt-Ca²⁺ uptake, Ca²⁺ buffering, and respiratory activity. Instead, it plays a conserved role in anabolic metabolism and stress response. These findings redefine MCUR1 as a foundational mitochondrial and metabolic regulator with potential implications for understanding and treating human disease.

### *MCUR1* loss-of-function variant in a patient with a rare neurological phenotype

We identified a 31-year-old male with a biallelic variant in *MCUR1*, presenting with an early-adult-onset neurological disorder beginning at 25 years of age and slow clinical progression (**Fig. 1A**). The patient exhibited impaired balance and vision, with bilateral internuclear ophthalmoplegia, vertical nystagmus, brisk deep tendon reflexes, bilateral Babinski sign, cerebellar ataxia, and pathological laughter. Magnetic resonance imaging (MRI) of the brain identified symmetric bilateral hyperintensities of the pyramidal tract in both the internal capsule and in the brainstem, as well as lesions affecting the medial longitudinal fasciculus (**Fig. 1B**). Magnetic resonance spectroscopy of the brain was unremarkable. Cell count analysis, and measurements of protein and lactate levels in the cerebrospinal fluid were normal. Electroneuromyography did not show signs of a disorder of the peripheral nervous system or muscle. Altogether, metabolic and histological examinations ruled out the diagnosis of a classical disorder of the mitochondrial oxidative phosphorylation system (OXPHOS), given that routine blood tests and photometric analyses of muscle biopsies showed normal levels of creatine kinase and lactate, cytochrome c oxidase (COX)/succinate dehydrogenase (SDH) staining, respiratory chain complex activity, and mitochondrial DNA sequence integrity.

**Figure 1.**
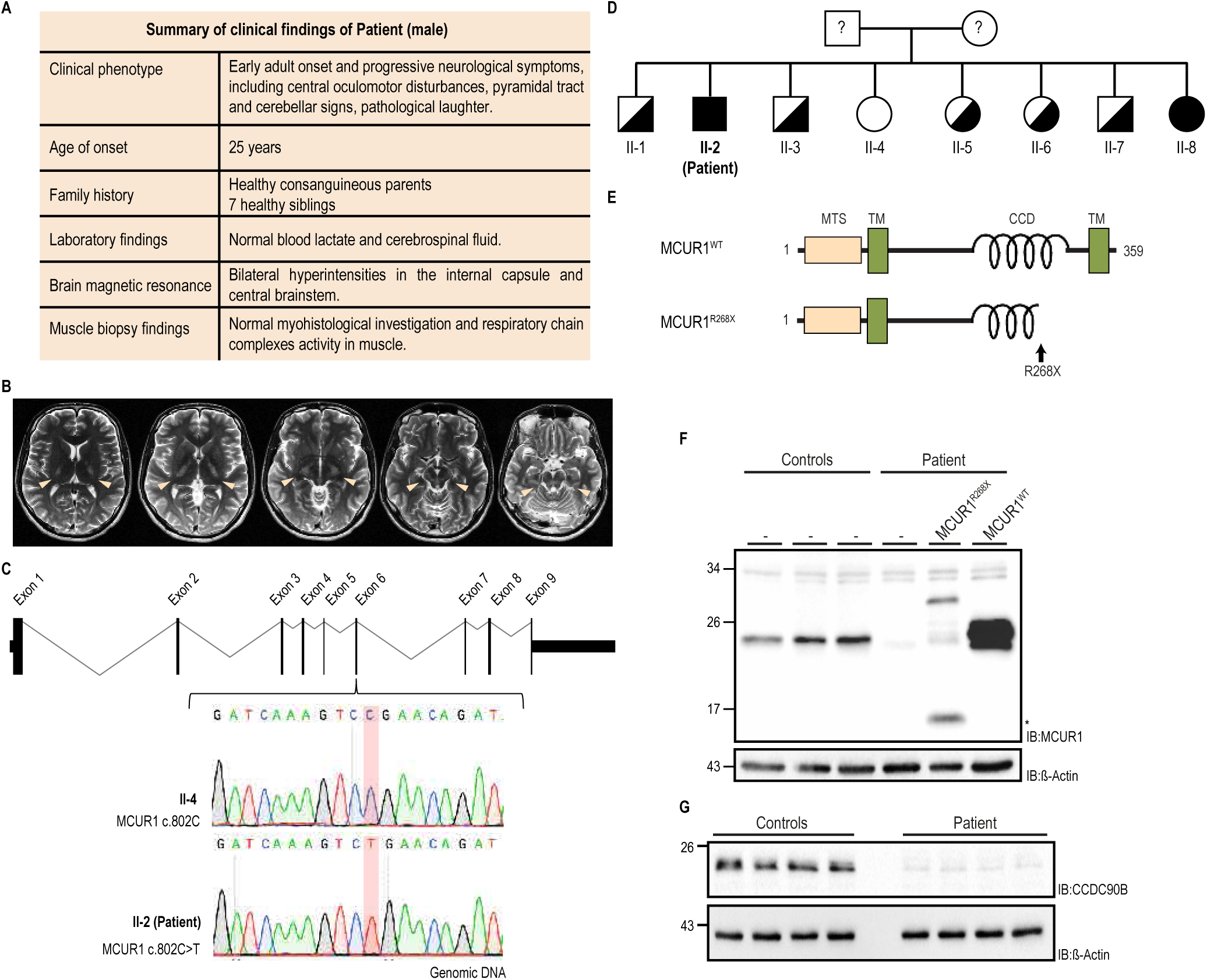
Identification of a *MCUR1* loss-of-function mutation in a consanguineous family. **(A)** Summary of clinical findings of the patient with *MCUR1* mutation. **(B)** Bilateral hyperintensities (beige arrow heads) of the internal capsule (*left*: pyramidal tract) and the medial longitudinal fasciculus (*right*: brain stem) identified by T2-weighted MRI scans of the patient’s brain. **(C)** Representation of *MCUR1* gene (ENSG00000050393), including its 9 exons. Whole-exome sequencing of patient (II-2)’s genomic DNA identified biallelic c.802C>T *MCUR1* variant in exon 6, as highlighted in red. Sequencing of the genomic DNA of sibling II-4 is represented as an example of non-mutated *MCUR1*. **(D)** Pedigree of the patient’s (II-2) consanguineous family. Males and females are represented by squares and circles, respectively, while filled and half-filled symbols refer to the family members being homozygous or heterozygous regarding the *MCUR1^c.802C>T^* variant, respectively. The patient (II-2) is marked in bold. The patient’s parents have not been sequenced so their genotype is unknown. **(E)** Protein domain representation of MCUR1 highlighting a mitochondrial targeting sequence (MTS), two transmembrane domains (TM), and a coiled-coil domain (CCD). The variant *MCUR1^c.802C>T^* is characterized by the introduction of an early stop codon on the amino acid R268 (represented with a black arrow). **(F)** Steady-state levels of MCUR1 in primary skin fibroblasts from three unrelated controls and the patient. Wild-type (WT) MCUR1 and MCUR1^R268X^ variants were expressed in fibroblasts derived from the patient. ß-actin is used as loading control. *, truncated variant of MCUR1. **(G)** Steady-state levels of CCDC90B in primary skin fibroblasts from one unrelated control and the patient (n=4 technical replicates). ß-actin is used as loading control. IB, immunoblot.

Whole-exome sequencing performed on total genomic DNA extracted from peripheral blood leukocytes identified a homozygous c.802C>T variant in *MCUR1* (ENST00000379170.9; rs367961059), which was confirmed by Sanger sequencing (**Fig. 1C**). As shown by the pedigree of the patient’s consanguineous family (**Fig. 1D**), five of the seven healthy siblings were heterozygous carriers of the same variant, whereas a 19-year-old sister (subject II-8) was a homozygous carrier but without any current clinical signs at the time of the investigation. This c.802C>T substitution occurs at low frequency in the Helmholtz Rare Disease Database (PhenoDis), in the Database for Single Nucleotide Polymorphism (dbSNP), and in the Genome Aggregation Database (gnomAD). In total, 50 heterozygous alleles were identified across these databases, which encompass sample sizes of 14,000, 56,250, and 1,607,006 individuals, respectively. The observed minor allele frequency of the c.802C>T variant is consistent with an autosomal recessive trait but may be more prevalent in ancestries underrepresented in current genomic datasets.

*MCUR1* encodes an integral protein of the inner mitochondrial membrane, containing two transmembrane domains and a coiled-coil domain. Both its N- and C-termini face the intermembrane space, while the intervening loop including the coiled-coil domain extends into the mitochondrial matrix (**Fig. 1E**). The biallelic c.802C>T *MCUR1* variant introduces a premature stop codon (R268X) within the coiled-coil domain of MCUR1, resulting in loss of function. Immunoblot analysis showed that the R268X substitution dramatically reduced MCUR1 protein levels compared with three unrelated control lines from unaffected individuals (**Fig. 1F**). By introducing an intron less wild-type MCUR1 (*MCUR1*^WT^), or the *MCUR1*^R268X^ isoform into patient skin fibroblasts, MCUR1 expression was restored, including the truncated MCUR1 protein. Thus, the low total MCUR1 expression levels observed in the patient results from nonsense-mediated mRNA decay triggered by the premature stop codon near the native exon-exon junction. Surprisingly, the protein levels of CCDC90B were also markedly reduced in the patient, suggesting a functional association between MCUR1 and CCDC90B (**Fig. 1G**).

### MCUR1 forms a stable hetero-oligomeric complex with CCDC90B

To elucidate the molecular function of MCUR1, we characterized its protein interaction network. We engineered a stable HEK-293 cell line expressing MCUR1 C-terminally tagged with StrepII-HA-His_6_ at near-endogenous levels, which served as bait for protein complex isolation (**Fig. 2A**). Subcellular fractionation (**Fig. S1A)**, carbonate extraction (**Fig. S1B**), and protease protection assays (**Fig. S1C**) confirmed correct mitochondrial targeting and membrane topology of the tagged protein. Tandem affinity purification (TAP) using C-terminal Strep II and His₆ tags, followed by LC-MS/MS analysis in five biological replicates with parental HEK-293 cells as negative controls, enabled discrimination of specific versus spurious interactions (**Fig. 2B**).

**Figure 2.**
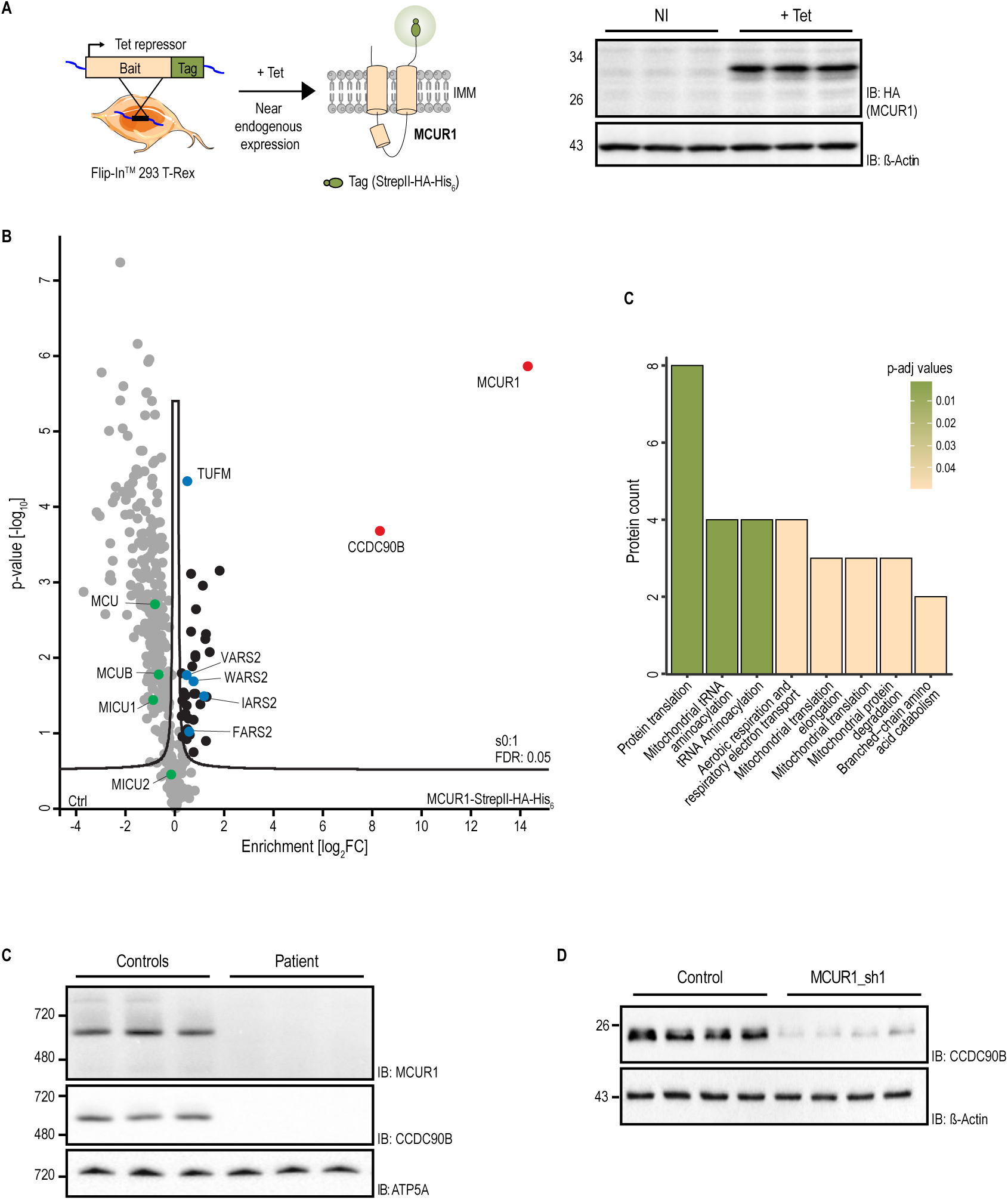
MCUR1 and its paralogue CCDC90B form a stable hetero-oligomeric complex. **(A)** Generation of Flp-In^TM^ 293 T-Rex cells expressing a C-terminally tagged MCUR1-StrepII-HA-His_6_ upon tetracycline-dependent induction. Whole-cell extracts before (-Tet) and after (+Tet) induction with tetracycline were separated by SDS-PAGE and analyzed by Western blotting. ß-actin is used as loading control. **(B)** Volcano plot of the mitochondrial interactome of MCUR1-StrepII-HA-His_6_ from lysates of tetracycline-induced Flp-In^TM^ 293 T-Rex cells.Cells not expressing the tagged protein are used as control (Ctrl). Possible binding partners were defined using a two-tailed Student t-test with a permutation-based false discovery rate (FDR) of 0.05 as a threshold. MCUR1 and CCDC90B are indicated in red; MCUC components are indicated in green. All significant hits are filled in black; blue dots highlight examples of mitochondrial protein translation-related factors that interact with the MCUR1–CCDC90B complex. **(C)** Pathway enrichment analysis (Reactome) for the 38 proteins enriched in the MCUR1-StrepII-HA-His_6_ pull-down highlights a connection between the MCUR1–CCDC90B complex and mitochondrial protein translation factors. **(D)** MCUR1 and CCDC90B form a protein complex absent in the patient with *MCUR1^c.802C>T^* variant. Blue Native-PAGE analysis of mitochondrial fractions from control and patient fibroblasts (*n*=3 biologically independent experiments). ATP5A was used as a loading control. **(E)** MCUR1 knockdown (MCUR1_sh1) in HeLa cells leads to loss of CCDC90B compared with control (pLKO) (n=4 technical replicates). ß-actin is used as loading control.

CCDC90B emerged as the most significantly enriched MCUR1 interactor, whereas components of the MCUC were not detected in association with MCUR1. Notably, previous studies that proposed MCUR1 as an MCUC subunit relied mainly on co-immunoprecipitation from overexpressed bait proteins. In contrast, unbiased proteomics surveys performed under near-endogenous conditions consistently failed to detect stable MCUR1–MCUC interactions^8^. Instead, our dataset revealed significant associations between MCUR1 and proteins involved in mitochondrial translation, including several aminoacyl-tRNA synthetases (e.g., VARS2, FARS2, WARS2, and IARS2) (**Fig. 2C**), many of which are linked to neurological disorders and brain abnormalities. Blue native (BN)-PAGE analysis of control fibroblast mitochondria identified a high-molecular weight complex containing both MCUR1 and CCDC90B, which was absent in patient fibroblasts (**Fig. 2D**). To control for genetic background effects, we generated isogenic HeLa cell lines expressing small hairpin RNAs (shRNAs) targeting MCUR1 (**Fig. S2**). Immunoblotting confirmed that MCUR1 and CCDC90B form a stable complex whose integrity depends on MCUR1 levels (**Fig. 2E**).

### MCUR1 deficiency does not impair mt-Ca^2+^ uptake or mitochondrial respiration

To evaluate whether the proposed role of MCUR1 in mt-Ca^2+^ regulation contributes to the patient phenotype, we quantified mt-Ca^2+^ transients using a matrix-targeted aequorin (mt-AEQ) probe in both intact and permeabilized patient fibroblasts. Upon histamine stimulation, which triggers Ca^2+^ release from the endoplasmic reticulum (ER), mt-Ca^2+^ uptake kinetics were indistinguishable between patient and control cells (**Fig. S3A**), and similar results were obtained in MCUR1 knockdown HeLa cells (**Fig. S3B**). To exclude potential compensatory changes in upstream Ca^2+^ signaling, we permeabilized the plasma membrane with digitonin upon depletion of intracellular Ca^2+^ stores by the SERCA inhibitor thapsigargin in presence of EGTA and triggered Ca^2+^ uptake with an exogenous Ca^2+^ bolus. mt-Ca^2+^ uptake kinetics remained comparable across conditions, regardless of whether mitochondria were energized with RCCI-(**Fig. S3C**) or RCCIV-specific substrates (**Fig. S3D**). Likewise, fluorescence-based measurements of Ca^2+^ clearance using the non-permeable dye Ca-Green-5N revealed no difference in the rate of signal decay (**Fig. S3E**) or overall mt-Ca^2+^ buffering capacity, indicating similar responses of the mitochondrial permeability transition pore to Ca²⁺ overload (**Fig. S3F**). The lack of an mt-Ca^2+^ uptake phenotype was not due to compensatory adaptations in the expression levels (**Fig. S4A,B**) or assembly (**Fig. S4C,D**) of MCUC components, indicating that MCUR1 is dispensable for mt-Ca^2+^ uptake under these conditions.

Given previous suggestions that MCUR1 affects RCC IV^5^ biogenesis, we next assessed mitochondrial bioenergetics in patient fibroblasts and HeLa MCUR1 knockdown cells. Oxygen consumption rates (OCR) analyses revealed no differences in basal, ADP-stimulated, or carbonyl cyanide m-chlorophenylhydrazone (CCCP)-uncoupled respiration between patient and control fibroblasts (**Fig. S5A-D**), while shMCUR1 HeLa cells showed only minor reductions (**Fig. S5E**). Expression and assembly of individual RCC subunits were also unaffected (**Fig. S6A-D**).

Finally, deletion of the MCUR1 homolog *FMP32* in two genetic strains of *Saccharomyces cerevisiae* did not impair growth on fermentable or non-fermentable carbon sources at any tested temperature (**Fig. S7A,B**). These results are consistent with previous findings^7,9^ and confirm that mitochondrial respiration remains intact in the absence of *FMP32*. Together, our data from human fibroblasts, HeLa cells, and yeast demonstrate that the primary role of MCUR1 extends beyond mt-Ca^2+^ handling and OXPHOS function, pointing instead to a broader role in mitochondrial metabolic regulation.

### Loss of MCUR1 and CCDC90B orthologs impairs nitrogen utilization and amino acid metabolism

To gain insights into MCUR1 function, we analyzed its conservation across 247 fully sequenced eukaryotic species (**Fig. S8**). MCUR1 homologs were identified in nearly all vertebrates and plants, but were absent from arthropods, apicomplexa (e.g., *Plasmodium falciparum*), and mitochondria-lacking protists (e.g., *Entamoeba histolytica*, *Giardia lambia*, *E. cuniculi*). Unlike MCUC components^10^, MCUR1 is broadly conserved in fungi, suggesting that its function extends beyond mt-Ca^2+^ uptake regulation. By contrast, CCDC90B exhibits a more restricted phylogenetic distribution, co-occurring with MCUR1 primarily in vertebrates, plants, and a few filamentous fungi—including the fission yeast *Schizosaccharomyces pombe* (*S. pombe*)—but not in *Saccharomyces cerevisiae* (**Fig. S8**). This distinct evolutionary pattern prompted us to investigate the conserved metabolic roles of MCUR1 and CCDC90B using *S. pombe*, which retains both homologs.

We identified three *S. pombe* sequences related to MCUR1: Sp-MCUR1 (*SPAC2C4.09*) and Sp-CCDC90B (*SPAC3H1.08c*), paralogous to human *MCUR1* and *CCDC90B*, respectively, and *SPBC27B12.07*, belonging to a distinct fungal clade (**Fig. 3A**). Given previous links between *S. cerevisiae* Fmp32 and amino acid utilization^7,9^, and between Sp-CCDC90B and regulation of the target of rapamycin complex 1 (TORC1)^11^, we examined whether *MCUR1* and *CCDC90B* contribute to nitrogen metabolism in *S. pombe.* Single (Sp-MCUR1 KO; Sp-CCDC90B KO) and double (Sp-double KO) deletion mutants in the prototrophic strain 972h, which can synthesize organic building blocks from inorganic sources, exhibited severe growth defects in minimal medium containing glucose and ammonium sulfate as the sole carbon and nitrogen sources. These defects were only partially rescued when proline replaced ammonium sulfate as the nitrogen source (**Fig. 3B**), implicating a role for MCUR1 and CCDC90B in amino acid metabolism. Moreover, wild-type and Sp-MCUR1 KO strains showed comparable growth on proline-derived metabolites such as arginine and glutamate (**Fig. S9A,B**), implicating a role for MCUR1 and CCDC90B in broader mitochondrial amino acid metabolism, rather than in the regulation or utilization of a single amino acid substrate. Importantly, the growth defect was rescued by combined supplementation with ammonium sulfate and glutamate (**Fig. S9C**), showing that upstream carbon/nitrogen supply can bypass the biosynthetic bottleneck caused by MCUR1–CCDC90B loss.

**Figure 3.**
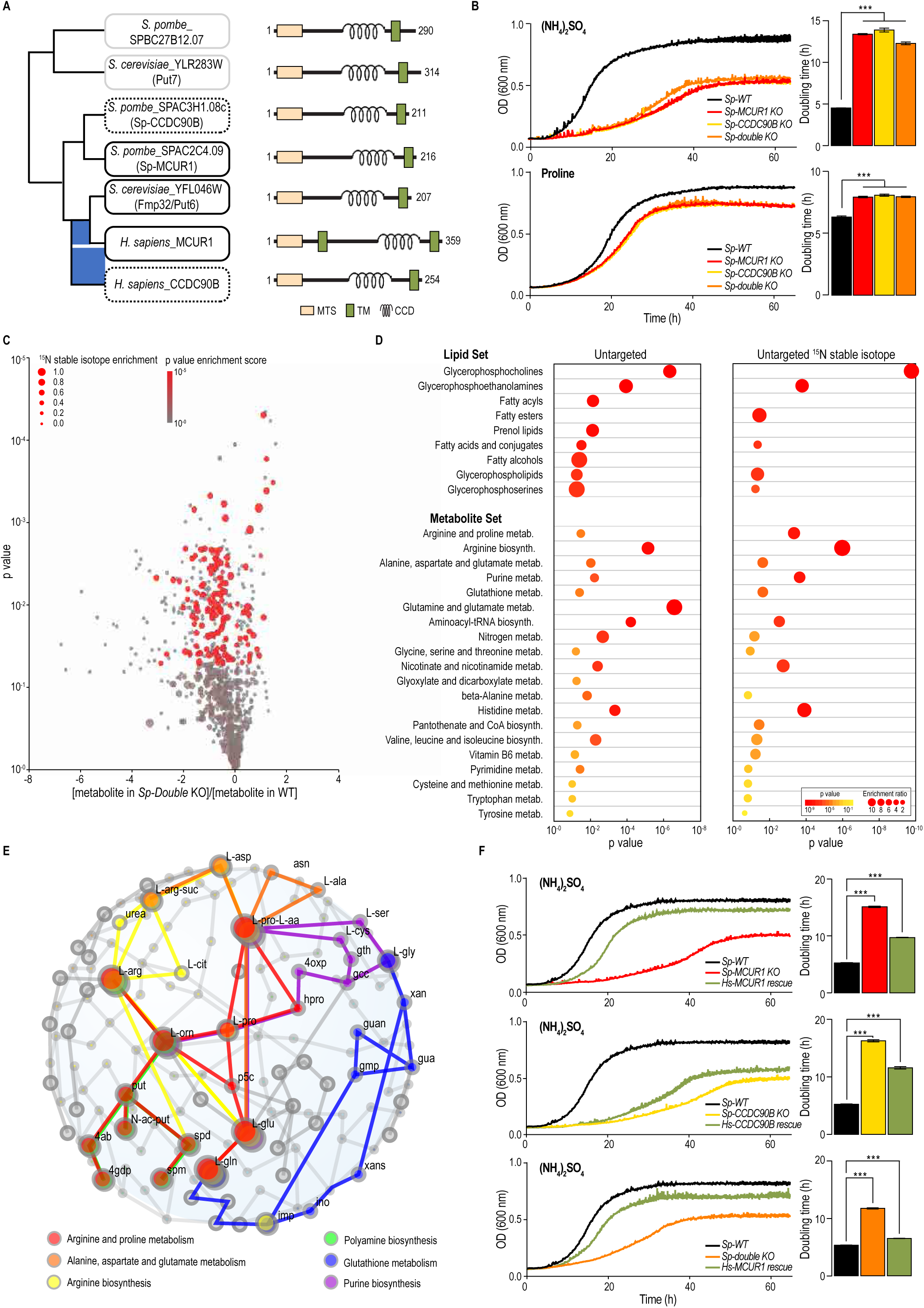
Impairment of amino acid metabolism by MCUR1 loss-of-function is conserved from yeast to human. **(A)** Phylogeny and protein features of MCUR1 and CCDC90B orthologs. Profile hidden Markov model (hmmsearch), using DUF1640 protein domain as key-word, retrieved three proteins of this family in *Schizosaccharomyces pombe* (*S. pombe*), including the orthologs of human MCUR1 (*SPAC2C4.09*) and CCDC90B (*SPAC3H1.08c*), here referred to as *Sp-MCUR1* and *Sp-CCDC90B*, respectively. MCUR1 orthologs are represented in black boxes and CCDC90B orthologs are represented in dotted black boxes. **(B)** Growth of prototrophic *S. pombe* wild-type (*Sp-WT*) and knockout strains lacking the homologs of either MCUR1 (*Sp-MCUR1 KO*), CCDC90B (*Sp-CCDC90B KO*), or both (*Sp-double KO)* in minimal media with either ammonium sulfate or proline as sole nitrogen source. Doubling time for each of the strains is represented on the right of the curves (*n*=4 biologically independent experiments; \*\*\**p* < 0.001, one-way ANOVA with Dunnett’s multiple comparisons test). **(C)** Untargeted stable isotope metabolomics analysis highlights enrichment of ^15^N stable isotope label in central carbon metabolites. Metabolomics profiling of WT versus Sp-MCUR1 and Sp-CCDC90B double knockout strains grown in minimal media with ammonium sulfate by untargeted (grey) and ^15^N stable isotope labeling LC-MS (red) with *p* < 0.05. **(D)** Pathway enrichment analysis of untargeted and ^15^N stable isotope labeled metabolites in *Sp-double KO*, represented in (C). Enrichment ratio of compound classes and metabolic pathways is depicted by sphere size, with significantly enriched sets highlighted in red with *p* < 0.05. **(E)** Metabolic network and pathway analysis of stable isotope enriched metabolomics in (C). Network perturbation of amino acid pathways is color coded highlighting interconnectedness of arginine and proline metabolism (red), alanine, aspartate and glutamate metabolism (orange), arginine metabolism (yellow), with polyamine biosynthesis (green), glutathione metabolism (blue), and purine biosynthesis (purple). **(F)** *Homo sapiens* MCUR1 (*Hs-MCUR1*) can functionally complement *S. pombe* single and double *Sp-MCUR1* and *Sp-CCDC90B* knockout strains and rescue their growth defect in minimal media supplemented with ammonium sulfate as the sole nitrogen source. Doubling time for each of the strains is represented on the right of the curves (*n*=4 biologically independent experiments; \*\*\**p* < 0.001, one-way ANOVA with Dunnett’s multiple comparisons test; all data represent mean ± SEM).

Untargeted stable-isotope-enhanced metabolomics using ^14^N/^15^N-labelled ammonium sulfate revealed widespread metabolic alterations in the Sp-double KO strain. After correcting for nitrogen-enhanced analytes and pathways, we observed pronounced changes in glycerophospholipids, fatty acid conjugates, and specific amino acid pathways including proline, arginine, alanine, aspartate, and glutamate (**Fig. 3C-E**), consistent with a broad multi-node metabolic dysregulation rather than a single-enzyme defect. Alterations in nicotinamide, purine, polyamine, and glutathione cycles pointed to redox and nitrogen balance disturbances with feedbacks to cellular turnover and stress responses. Expression of human MCUR1–but not human CCDC90B–partially restored growth in both single and double mutants (**Fig. 3F**), demonstrating that MCUR1 has a conserved role in maintaining mitochondrial amino acid homeostasis and exerts a dominant functional effect over CCDC90B.

### MCUR1 deficiency in humans impairs amino acid metabolism

To evaluate the systemic consequences of MCUR1 deficiency, we performed untargeted metabolomic profiling of patient urine and serum and compared metabolite levels to reference datasets representing normal physiological conditions, including fasting, dietary intake, physical activity, and standardized metabolic challenge tests (**Fig. 4A,B**). Arginine, hydroxyproline, carnitine, taurine, valine, and isoleucine were significantly accumulated (*p* < 0.05), and pathway enrichment analyses revealed perturbations in arginine and proline metabolism, glycine and serine metabolism, taurine metabolism, alongside dysregulation of nitrogen, gluthathione, and tricarboxylic acid (TCA) cycle (**Fig. 4A,B**). Aminoacyl-tRNA biosynthesis, a process encompassing enzymes that interact with MCUR1-CCDC90B complex (**Fig. 2B**), was also among the enriched pathways. Despite a life-long vegetarian diet, the patient displayed a marked accumulation of biomarker metabolites typically associated with a carnivorous diet. Additional dysregulations were detected in the urea cycle, ammonia recycling, glutamate, aspartate, and glutathione metabolism, as well as in spermidine/spermine biosynthesis, branched-chain amino acids (valine, leucine, and isoleucine), and the TCA cycle (*p<* 0.05). Complementary whole-cell proteomics of patient-derived fibroblasts and MCUR1 knockdown HeLa cells (**Fig. 4C** and **Fig. S9D**) revealed a consistent upregulation of vacuolar, phagosomal, and lysosomal proteins, together with activation of autophagy-related pathways, oxidative stress, and energy metabolism, accompanied by downregulation of cell cycle regulators, E2F target proteins, epithelial-mesenchymal transition (EMT) processes, RNA processing factors, and mitochondrial metabolic enzymes, indicating a coordinated impairment in biosynthetic and proliferative capacity (**Fig. 4D**). Collectively, these multi-omics signatures collectively point toward a bottleneck in the production or utilization of non-essential amino acids and lipids in MCUR1-deficient cells—pathways that critically depend on metabolic shuttles across the mitochondrial membrane but could be partially compensated via anaplerotic amino acid salvage routes.

**Figure 4.**
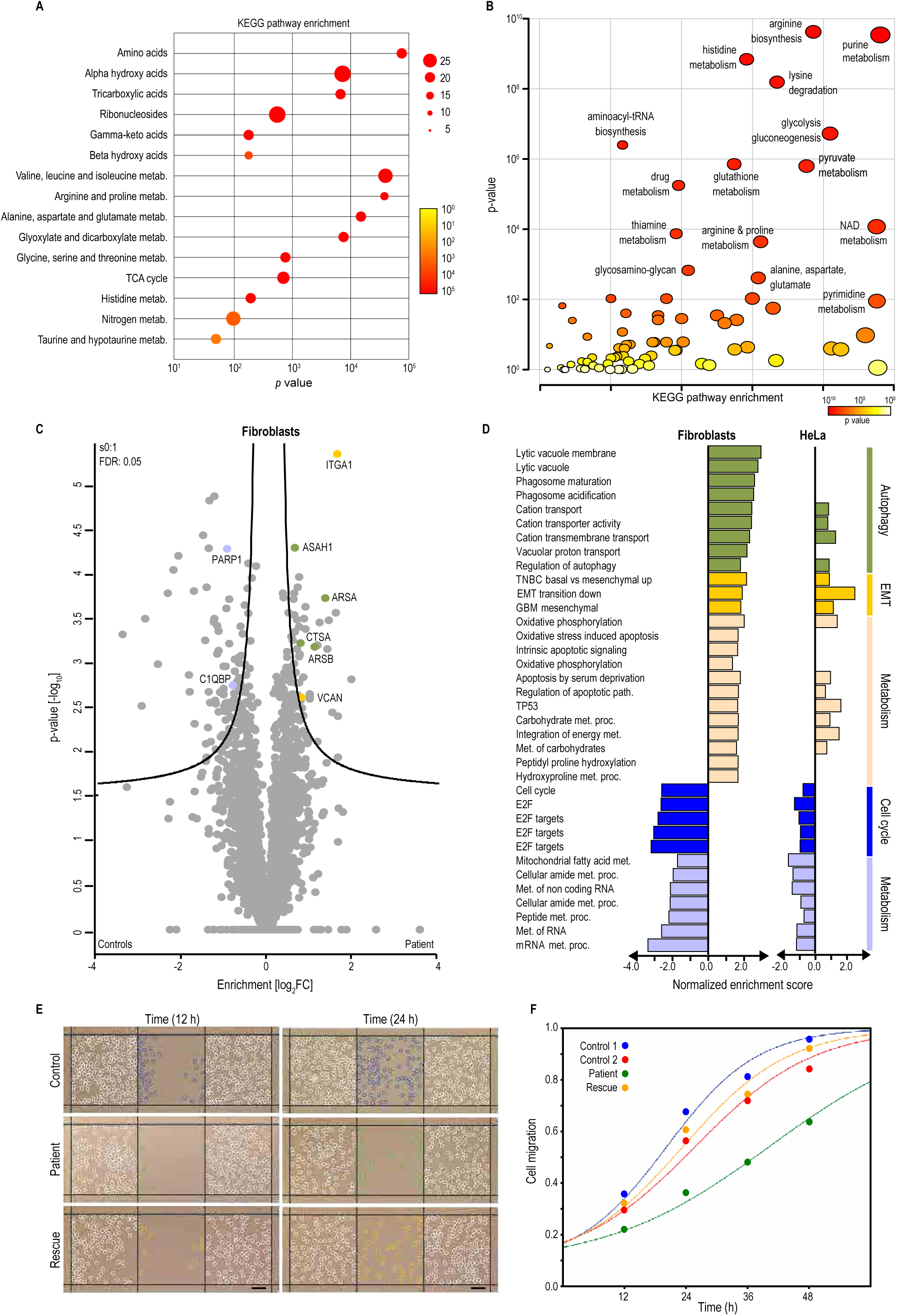
Multi-omics profiling identifies a dysregulation of amino acid metabolism in the MCUR1 patient. **(A)** Metabolic profiling and pathway enrichment analysis of urine samples from patient and healthy controls. Healthy controls urine data refer to the HuMet repository^25^. Enrichment ratio of compound classes and metabolic pathways is depicted by sphere size, with significantly enriched sets highlighted in red with *p* < 0.05. **(B)** Pathway enrichment analysis of untargeted serum metabolomics from patient in comparison to healthy controls (*p* < 0.05). Healthy controls serum data refer to the HuMet repository^25^. **(C)** Proteomics profiling in human fibroblasts from control individuals and the patient (II-2). The Volcano plot shows significant differently expressed proteins (filled black circles) between control and patient fibroblasts (*p* values< 0.05). CCDC90B was identified only in the control samples (FDR 0.05, S0 0.1). Examples of proteins related to vacuole, EMT transition and RNA metabolism are marked in yellow, green and blue, respectively. **(D)** Pathway enrichment analysis of whole cell proteomics from (C) and Fig. S9E. Pathways with positive normalized enrichment score in MCUR1 deficient proteomes are colored in green-to-beige shades, while downregulated ones are indicated in blue shades (*p* < 0.05 and *q* < 0.1). **(E)** Cell migration scratch wound healing assay. Controls, patient-derived fibroblasts, and MCUR1 wild-type rescue of patient-derived fibroblasts were grown in minimal essential media supplemented with serum and uridine after scratch generation. Cell migration was monitored by bright light microscopy in combination with digital image acquisition and data quantification at 0, 12, 24, 36, and 48 hours. **(G)** Quantification of cell migration in cells grown as in (E). Data represent mean ± SD (*n*=3 biologically independent experiments).

To assess the functional impact of this metabolic defect, we performed *in vitro* wound-healing assays using control and MCUR1-deficient fibroblasts cultured in minimal media lacking non-essential amino acids. Patient fibroblasts displayed markedly delayed growth and migration compared with controls, a phenotype that was fully rescued by WT MCUR1 re-expression (**Fig. 4E,F** and **Fig. S9E**), confirming a causal role for MCUR1 in supporting metabolism for sustaining cell proliferation. Together, these data demonstrate that *MCUR1* deficiency compromises non-essential amino acid metabolism, limits biosynthetic capacity, and impairs cell proliferation, particularly under nutrient-restrictive conditions.

In summary, by integrating biochemical, proteomic, and evolutionary analyses, our study reveals that the MCUR1–CCDC90B complex functions as a conserved mitochondrial scaffold coupling nitrogen metabolism, amino acid and lipid shuttling, and translational capacity to cellular growth control. MCUR1 forms a stable complex with CCDC90B, distinct from the canonical MCU calcium uniporter, and interacts with mitochondrial transporters and aminoacyl-tRNA synthetases that link protein synthesis to metabolic flux. Across human cells, yeast, and patient-derived fibroblasts, loss of MCUR1 impairs utilization of inorganic nitrogen sources, disrupts mitochondrial homeostasis without altering MCU complex formation, and causes vacuolar and autophagy-related dysfunction. Metabolically, MCUR1 deficiency leads to dysregulation of amino acid intermediates and lipid species, reflecting a broad disruption of carbon flux connecting the amino acids glutamate, glutamine, ornithine, proline, and hydroxyl-proline with adjacent TCA cycle-related intermediates—hallmarks of a metabolic bottleneck in mitochondrial non-essential amino acid metabolism. Functionally, this metabolic defect constrains cell migration and growth under nutrient limitation, positioning MCUR1 as a critical regulator of mitochondrial metabolic capacity. These findings establish a mechanistic framework linking mitochondrial metabolic integrity to neurological health and identify the MCUR1–CCDC90B module as a central node coupling mitochondrial metabolism to organellar, cellular, and organismal homeostasis. Consistent with this pleiotropic role, MCUR1 is highly expressed in energy-demanding organs such as the brain, liver, and heart^12^, which rely on amino acid and lipid metabolism to sustain biosynthetic and redox balance. The brain, in particular, depends on glutamate, glutamine, and branched-chain amino acids for neurotransmitter synthesis and neuronal signaling, providing a plausible link between MCUR1 dysfunction and the observed neuropathological phenotype. Finally, developmental, dietary, or epigenetic factors may modulate disease penetrance and onset, as illustrated by the asymptomatic younger homozygous sibling. Future work should delineate how the MCUR1–CCDC90B complex coordinates amino acid and lipid metabolism with stress responses and autophagy, and whether these biochemical perturbations directly underlie the neurological manifestations in affected patients.

## METHODS

### Genetic testing

After obtaining patient consent, total genomic DNA was extracted from peripheral blood leukocytes of the patient and subjected to whole exome analysis. Coding regions were enriched using a SureSelect Human All Exon V5 kit (Agilent) followed by sequencing as 100-bp paired-end runs on an Illumina HiSeq2500. Reads were aligned to the human reference genome (UCSC Genome Browser build hg19) using Burrows-Wheeler Aligner (v.0.7.5a) and single-nucleotide variants and small insertions and deletions (indels) were detected with SAMtools (version 0.1.19). For segregation analysis Sanger sequencing was performed in all siblings using DNA extracted from peripheral blood leukocytes or buccal epithelial cells.

### Cell lines

All mammalian cells were grown in high-glucose Dulbecco’s modified Eagle’s medium (DMEM) supplemented with 10% FBS at 37°C and 5% CO_2_. HeLa cells stably expressing a wild-type mitochondrial matrix-targeted GFP-aequorin (mt-AEQ) were grown in presence of 100 μg/ml geneticin^13^. Primary human skin fibroblasts were grown in presence of 0.2 mM uridine. Primary skin fibroblasts stably expressing wild-type human MCUR1 and mt-AEQ from the pLX304 lentiviral vector were generated by transduction, as previously described^14^, and grown in presence of 0.2 mM uridine and 10 ug/mL blasticidin. HeLa mt-AEQ cells stably expressing either the lentiviral empty vector pLKO or a pLKO vector expressing sh-RNAs targeting human MCUR1 (sh1, TRCN0000134711; sh2, TRCN0000134319; sh3, TRCN0000135627; sh4, TRCN0000136480; sh5, TCRN0000137294) were generated as previously described^15^ and grown in presence of 2 ug/mL puromycin. Lentivirus production and infection were performed according to guidelines from the Broad RNAi Consortium and infected cell lines were selected 48 hr post-transduction with the respective selection markers. Primary control skin fibroblasts were kindly provided Prof. Vincent Bonifati (Erasmus University Medical Center). Parental Flp-In^TM^ 293 T-Rex cells were cultured in presence of 15 µg/mL blasticidin and 100 µg/mL Zeozin. Flp-In^TM^ 293 T-Rex cells stably expressing a C-terminal StrepII_2_-HA-His_6_-tagged MCUR1 under the control of a tetracycline-regulated CMV/TetO_2_ promoter were grown in presence of 15ug/mL blasticidin and 100ug/mL hygromycin.

### SDS/BN-PAGE and immunoblotting

Immunoblot analyses were performed with the following antibodies: MCU (Sigma Aldrich, HPA01648), anti-ATP5A (Abcam, MS507), NDUFB8 (Abcam, ab110242), SDHA (Abcam, ab14715), UQCRC2 (Abcam, ab14745), MTCO II (Abcam, ab110258), EMRE (Santa Cruz, sc-86337), MCUB (Santa Cruz, sc-163985), MCUR1 (Thermo Fisher, PA5-32164), CCDC90B (Sigma Aldrich, HPA011931),TIM23 (BD Bioscience, 611222), TOM20 (Abcam, ab56783), Cyclophilin D (Abcam, ab110324), MICU1 (Atlas Antibody, HPA037479), MICU2 (Sigma Aldrich, HPA045511), Actin (Sigma Aldrich, A2228), HA (BioLegend, MMS-101R), VDAC1/Porin (Abcam, ab14734), Lamin A/C (abcam, ab238303). BN-PAGE was performed as previously described^14^. Briefly, crude mitochondria were solubilized for 10 min on ice in 1xNativePAGE^TM^ sample buffer containing 1% (w/v) digitonin and after a centrifugation step at 20,000 x *g* for 30 min at 4°C, the supernatant was mixed with 0.25% of NativePAGE^TM^ G-250 sample additive and resolved on a 3-12% gradient NativePAGE^TM^ Novex Bis-Tris gel according to the manufacturer‘s protocol. Proteins were transferred onto PVDF membranes by electrophoretic wet transfer overnight at 40V, 4°C. After transfer, proteins were fixed on the membrane by incubating in 8% acetic acid for 15 min at room temperature for immunoblot analyses.

### Mitochondrial bioenergetics in intact and permeabilized cells

Glycolytic and mitochondrial respiratory activities were assessed by microplate-based respirometry using a Seahorse Extracellular Flux (XF) Analyzer (Agilent Technologies, Santa Clara, USA) as previously described^16^. Briefly, cells were seeded on a Seahorse XF cell culture 96-well plate in growth media and incubated overnight at 37°C and 5% CO_2_. For measurements of oxygen consumption rate (OCR) and extracellular acidification rate (ECAR) in intact cells, the growth medium was replaced with either mito-stress assay (MSA) medium containing pyruvate (1 mM), L-glutamine (2 mM), and glucose (10 mM) or glycolysis assay medium with L-glutamine (2mM). Cells were then incubated at 37°C and 5% CO_2_ for 1 hour. For measurements of For OCR measurements in permeabilized cells, cells were washed with PBS and incubated for 5 min in a mito-assay medium (MAS) containing d-mannitol (200 mM), d-sucrose (70 mM), 2.5 MgCl_2_ (5 mM), KH_2_PO_4_ (10mM), HEPES-KOH (2 mM), EGTA-KOH (1 mM), 0.5% fatty-acid free BSA, and digitonin (60 µM for HeLa cells and 25 µM for primary skin fibroblasts) to selectively permeabilize the plasma membrane. Cells were then washed with MAS and OCR was measured in MAS containing respiratory substrates. OCR and ECAR measurements were normalized to cell number by quantifying total DNA with the CyQUANT Cell Proliferation Assay Kit.

### Isolation of crude mitochondria from cultured cells

Crude mitochondria were isolated from cultured cells as previously described^14^. Briefly, cells were grown to confluency, washed three times with PBS, scraped down and resuspended in PBS. After 5 min of centrifugation at 600 x *g*, 4°C, the cell pellet was resuspended in ice cold isolation buffer (IB: 220 mM d-mannitol, 70 mM d-sucrose, 5 mM HEPES-KOH pH 7.4, 1 mM EGTA-KOH pH 7.4, 0.5% fatty acid-free BSA, protease inhibitors). The plasma membrane was permeabilized by nitrogen cavitation at 600 psi for 10 min at 4°C. The whole cell homogenate was then centrifuged at 600 x *g* for 10 min at 4°C and the pellet (including the nuclei fraction) was saved for immunoblotting. The supernatant was transferred into new tubes and centrifuged at 8000 x *g* for 10 min at 4°C. The supernatant (corresponding to the cytosolic fraction) was saved for immunoblotting and the resulting pellet containing crude mitochondria was resuspended in IB for further analyses.

### Measurements of mitochondrial calcium uptake in intact human cells

Mitochondrial Ca^2+^ uptake was measured in intact HeLa cells and primary skin fibroblasts expressing mt-AEQ as previously described^13^. Briefly, cells were seeded in white 96-well plates in growth medium without selection marker and after 24 hours, mt-AEQ was reconstituted with 2 uM native coelenterazine for 2 hours at 37°C and 5% CO_2_. AEQ-based, Ca^2+^-dependent light kinetics were measured upon stimulation with histamine (100 uM). Light emission was measured in a luminescence counter (MicroBeta2 LumiJET Microplate Counter, PerkinElmer) at 469 nm every 0.1 s. At the end of each experiment, cells were lysed with a solution containing 0.5% Triton X-100 and 10 mM CaCl_2_ to release all the residual aequorin counts. Quantification of mt-Ca^2+^ concentration was performed as described also as previously described.

### Measurements of mitochondrial calcium uptake in digitonin-permeabilized human cells

Mitochondrial Ca^2+^ uptake was measured in permeabilized HeLa cells and primary skin fibroblasts expressing mt-AEQ as previously described^14^. Briefly, cells were harvested at a density of 500,000 cells/mL in growth medium containing 20 mM HEPES-NaOH, pH 7.4. AEQ was reconstituted with 3 uM native coelenterazine for 2 hours at room temperature and the cell pellet was then incubated in an extracellular-like buffer (145 mM NaCl, 5 mM KCl, 1 mM MgCl_2_, 10 mM glucose, 10mM HEPES and 500 uM EGTA pH 7.4/NaOH) containing 200 nM thapsigargin for 20 min at room temperature. Cells were re-suspended in an intracellular-like buffer (140 mM KCl, 1 mM KH_2_PO_4_/K_2_HPO_4_, 1 mM MgCl_2_, 1 mM ATP/MgCl_2_, 20 mM HEPES and 100 uM EGTA pH 7.2/KOH) supplemented with respiratory substrates. Cells were permeabilized with either 60 uM (HeLa cells) or 25 μM (human skin fibroblasts) digitonin for 5 min and re-suspended in intracellular-like buffer at a density of 900 cells/uL. Then, 90 uL of cell suspension was dispensed into a white 96-well plate and Ca^2+^-stimulated light signal was recorded at 469 nm every 0.1 s using a luminescence counter (MicroBeta2 LumiJET Microplate Counter, PerkinElmer). At the end of each experiment, cells were lysed with a solution containing 0.5% Triton X-100 and 10 mM CaCl_2_ to release all the residual aequorin counts. Quantification of mt-Ca^2+^ concentration was performed as described in Arduino et al. (2017). Measurement of mt-Ca^2+^ buffering capacity in digitonin permeabilized human cells was performed as previously described using Calcium Green-5N Hexapotassium Salt, cell impermeant. Calcium Green-5N fluorescence (excitation 506 nm, emission 531 nm) was monitored every 2 s at room temperature using a CLARIOstar microplate reader (BMG Labtech) upon injection of CaCl_2_. The MCU inhibitor Ru360 (10 uM) was used as a positive control.

### Stable protein expression in Flp-In^TM^ 293 T-Rex cells

Flp-In^TM^ 293 T-Rex cells stably expressing a C-terminal StrepII_2_-HA-His_6_-tagged MCUR1 under the control of a tetracycline-regulated CMV/TetO_2_ promoter were generated as previously described^8^. First, the open reading frame (ORF) of MCUR1 was cloned into the pcDNA5/FRT/TO Flp-In expression vector, in frame with a C-terminal StrepII_2_-HA-His_6_-tag. Flp-In^TM^ 293 T-Rex cells contain a single integrated flippase (Flp) recognition (FRT) site and stably express the *lacZ-ZeocinTM* fusion gene (pFRT/*lac*Zeo) and the tetracycline repressor (pcDNA6/TR). Co-transfection of Flp-In^TM^ 293 T-Rex cells with the Flp-In^TM^ expression vector and the Flp recombinase vector pOG44 results in a targeted integration of MCUR1 at a single transcriptionally active genomic locus that is the same in each cell. Briefly, cells were seeded in 6-well plates without antibiotics and transfected with 80 ng of the Flp-In™ expression vector and 720 ng of pOG44 using X-tremeGENE^TM^ HP DNA according to manufacturer’s protocol. Stable transfectants were selected and maintained in growth medium containing blasticidin (15ug/mL) and hygromycin (100ug/mL). The expression of the tagged bait protein was induced with 1 µg/mL tetracycline in media without antibiotics. After 24 hours, cells were washed three times with PBS and the cell pellet was collected at 600 x *g* for 10 min at 4°C and stored at -80°C till further use.

### Tandem affinity purification

A cell pellet containing ∼90 x 10^6^ cells was re-suspended in 2 mL lysis buffer (50mM HEPES-KOH pH7.4, 150mM KCL, 5mM EGTA, 5% glycerol, 3% digitonin, protease inhibitors), incubated for 30 min at 4 °C on a rotating shaker, and centrifuged at 10 000 x *g* for 5 min at 4 °C. The supernatant was mixed with HEPES-KOH pH 8 (12.5 mM) and avidin (10 uM) and incubated for 20 min at room temperature on a rotating shaker. The resin (150 uL streptavidin) was washed three times with 500 µL washing buffer (WB: 50 mM HEPES-KOH pH 7.4, 150 mM KCl, 5 mM EGTA, 5% glycerol, 0.02% digitonin), incubated with the supernatant for 45 min at 4 °C on a rotating shaker, and washed again three times with WB. Next, the resin was incubated with 250 µL of elution buffer (EB: 50 mM HEPES-KOH pH 8, 150 mM KCl, 5 mM EGTA, 5% glycerol, 10 mM biotin) for 5 min at room temperature and the eluate was collected with a quick spin. The same procedure was repeated three times and the eluates were then combined and incubated with 50 uL of Ni-NTA resin for 45 min at 4 °C on a rotating shaker. Any unbound fraction was removed by washing three times with 500 µL WB. For SDS-PAGE analysis, the resin was resuspended in 1x Laemmli, whereas for LC-MS/MS analysis the resin was washed three times with digitonin-free WB followed by incubation with 35 µL RapiGest for 10 min at room temperature. Five volumes of ice-cold acetone were added to the eluate, the sample was vortexed thoroughly for 10 sec and incubated overnight at -20°C. The sample was centrifuged at 20 000 x g for 30 min at 4 °C and the pellet was stored at -80 °C for LC-MS/MS analysis.

### Sample preparation for mass spectrometry analysis

Eluates were resuspended in 50 µl denaturation buffer (6 M urea, 2 M thiourea, 10 mM HEPES pH 8.0, 10 mM DTT) for 30 min at RT before adding alkylation agent (55 mM iodoacetamide) and incubating at RT in the dark for 20 min. Proteins were digested at RT for 3 hrs by adding LysC at 1:100 ratio of enzyme:protein, before diluting the sample with 50 mM ammonium bicarbonate to reach urea concentration of 2M. Subsequently, samples were digested at RT overnight by adding Trypsin at 1:100 ratio of enzyme:protein. The next day, the digestion was stopped by acidifying the sample with 10 µL of 10% TFA and the final peptides were cleaned up using SDB-RPS StageTips as described in^17^.

### Whole cell proteome analysis

Whole proteome of Hela cells and human skin fibroblasts was analyzed by LC-MS/MS mass spectrometry. The cells were lysed by scraping them in a lysis buffer containing 1% sodium deoxycholate, 10 mM of the reducing agent tris(2-carboxyethyl)phosphine (TCEP) and 40 mM of the alkylating agent 2-chloroacetamide (CAA),chloroacetamide, an optimized reaction mix that allows reduction and alkylation to be performed as a single step (Kulak et al 2014). The lysate was boiled and sonicated using a Bioruptor system (10 cycles, 30 sec on/off; Diagenode, Belgium), acetone precipitated and then digested as described above. Samples (20 ug) were digested overnight with LysC and Trypsin in a 1:40 ratio and desalted using SDB-RPS StageTips.

### LC-MS/MS analysis

MS analysis was performed using Q Exactive HF mass spectrometers (Thermo Fisher Scientific, Bremen, Germany) coupled on-line to a nanoflow UHPLC instrument (Easy1000 nLC, Thermo Fisher Scientific). Peptides were separated on a 50 cm long (75 μm inner diameter) column packed in-house with ReproSil-Pur C18-AQ 1.9 μm resin (Dr. Maisch GmbH, Ammerbuch, Germany). Column temperature was kept at 50 °C. Peptides were loaded with buffer A (0.1% (v/v) formic acid) and eluted with a nonlinear gradient of 5-60% buffer B (0.1% (v/v) formic acid, 80% (v/v) acetonitrile) at a flow rate of 250 nl/min. Peptide separation was achieved by 120 min gradients for both whole proteomes and pulldowns. For pulldowns, the survey scans (300-1650 m/z, target value = 3E6, maximum ion injection times = 20 ms) were acquired at a resolution of 60,000 followed by higher-energy collisional dissociation (HCD) based fragmentation (normalized collision energy = 27) of up to 15 dynamically chosen, most abundant precursor ions. The MS/MS scans were acquired at a resolution of 15000 (target value = 1E5, maximum ion injection times = 60 ms). Repeated sequencing of peptides was minimized by excluding the selected peptide candidates for 20 seconds. For whole proteome analysis the survey scans were acquired at a resolution of 120,000, the MS/MS scans at a resolution of 15000 and a dynamic exclusion of 30 seconds.

### MS data processing

Interactome data were analysed using the MaxQuant software package 1.5.5.2^18^. The false discovery rate (FDR) cut-off was set to 1% for protein and peptide spectrum matches. Peptides were required to have a minimum length of 7 amino acids and a maximum mass of 4600 Da. MaxQuant was used to score fragmentation scans for identification based on a search with an initial allowed mass deviation of the precursor ion of a maximum of 4.5 ppm after time-dependent mass calibration. The allowed fragment mass deviation was 20 ppm. Fragmentation spectra were identified using the UniprotKB *Homo sapiens* database with the Andromeda search engine^19^. Carbamidomethylation of cysteine was set as fixed modification and N-terminal protein acetylation as well as methionine oxidation as variable modifications. Both ‘label-free quantification (MaxLFQ)’ with a minimum ratio count of 1 and ‘match between runs’ with standard settings were enabled^20^. Fibroblasts and HeLa proteome data were analysed using the MaxQuant version 1.6.0.12 with the same parameters. The mass spectrometric data have been deposited via PRIDE^21^ to the ProteomeXchange Consortium (http://proteomecentral.proteomexchange.org) under the accession number PXD017460.

### MS data analysis and visualization

Basic data handling, normalization, statistics, and annotation enrichment analysis were performed with the Perseus software package^22^. To generate the list of interactors of the three baits analyzed, we used the MaxLFQ, MaxQuant’s label-free quantification (LFQ) algorithm^20^. ProteinGroups were filtered removing hits to the reverse decoy database and proteins only identified by modified peptides. Differentially expressed proteins were identified by two-tailed Student t-test using the volcano plot option of Perseus.

### Analysis of mitochondrial protein topology

The topology of mitochondrial proteins was assessed by proteinase K (PK) protection assay on freshly isolated crude mitochondria as previously described^14^. Briefly, mitochondria were incubated for 15 min at room temperature in IB buffer containing increasing concentrations of digitonin or 1% TritonX-100 and 100ug/ml PK. PK was inactivated by incubation with 5 mM PMSF on ice for 10 min. Samples were then mixed with Laemmli buffer containing 10% 2-mercaptoethanol for immunoblot analysis. Alkaline carbonate extraction was performed by incubating crude mitochondria with 0.1 M Na_2_CO_3_ at pH 10 and pH 11 for 30 min on ice. Samples were then centrifuged at 45,000 x g for 10 min at 4°C and the pellets were resuspended in Laemmli buffer containing 10% 2-mercaptoethanol for immunoblot analysis. Supernatants were mixed with 100% TCA, incubated overnight at -20°C, and then centrifuged at 16,000 x g for 25 min at 4°C. Pellets were then washed twice with cold acetone, air-dried for 30 min at room temperature and resuspended in Laemmli buffer for immunoblot analysis.

### *Saccharomyces cerevisiae* strains and growth measurements

*FMP32* was deleted in BY4742 and YPH499 wild-type *S. cerevisiae* strains by homologous recombination using hygromycin or kanamycin cassettes. Yeast transformation was performed by lithium acetate method and transformants were selected on YPD (1% yeast extract, 2% peptone, and 2% glucose) plates containing either hygromycin (200µg/mL) or G418 (500µg/mL), respectively. Successful deletion of *FMP32* in individual clones was confirmed by isolation of genomic DNA and subsequent PCR using the following primer pairs: SacFmp32p (5’-CCC GAG CTC TT ACC GTC TCA ATT TCA CAA G -3’) and KanB (5’-CTG CAG CGA GGA GCC GTA AT -3’) and SacFpm32p and Fmp32Stop132Eco (5’-CCC GAA TTC CTA TTT GGT AAT AAC AAC CCT TAG – 3’). The former primer pair was used to confirm the successful deletion and the latter to exclude the presence of the wild-type copy of the gene. Genomic DNAs isolated from wild-type BY4742 strain and *Dfmp32:KAN* obtained from from the Saccharomyces Genome Deletion Project (http://www-sequence.stanford.edu/group/yeast deletion project/deletions3.html) served as controls for PCR reactions. For growth assays in liquid media, yeast cultures were grown at 30°C overnight in YPD media, diluted to an OD of 0.1 and then grown in black, gas-permeable Lumox 96-well plates. Absorbance was measured at 600 nm and at intervals of 340 s using a CLARIOstar microplate reader (BMG Labtech) for 24 hours at 30°C. For growth assays on agar plates, yeast cultures were grown at 30°C overnight in YPGal media (1% yeast extract, 2% peptone, and 2% galactose), diluted in the morning to an OD of 0.1 in fresh YPGal medium and grown at 30°C till they reached an OD of 0.5. Cells were harvested, washed with water and spotted on agar plates containing different carbon sources (YPD; YPG, 3% glycerol; YPL, 2% lactic acid) following a 10-fold serial dilution and incubated at 24 °C, 30 °C or 37 °C.

### *Schizosaccharomyces pombe* strains and growth measurements

Single and double knockout of SPAC2C4.09 (Sp-MCUR1) and SPAC3H1.08c (Sp-CCDC90B) in the prototroph *S. pombe* 972 h strain were constructed by homologous recombination by lithium acetate transformation of PCR products containing 500 bp of targeting homology using *KanMX6* and natMX6, respectively, as selection markers as described previously^23^. For expression of human MCUR1, CCDC90B, or both, codon-optimized sequences were integrated into the chromosomal loci of SPAC2C4.09 and SPAC3H1.08c and expressed under the control from their endogenous promoters, respectively. Insertions of *MCUR1* and *CCDC90B* were generated by replacing the deletion cassettes SPAC2C4.09::KanMX6 and SPAC3H1.08c::natMX6 individually with *MCUR1*::hygMX and *CCDC90B*::kanMX by homologous recombination. The correct genomic integration and expression of the human homologs were confirmed by diagnostics PCR using genomic DNA and quantitative PCR of total RNA with gene-specific primers (RT-qPCR), respectively. Yeast extract with supplements (YES) media containing 0.5 % (w/v) yeast extract, 3.0 % (w/v) glucose, 225 mg/l adenine, 225 mg/l histidine, 225 mg/l leucine, 225 mg/l uracil, 225 mg/l lysine hydrochloride) was used for liquid culture, unless specified otherwise. Deletion strains were selected on YES plates containing 100 mg/l G418, 100 mg/ hygromycin, and 50 mg/ml nourseothricin, respectively. Liquid growth assays were conducted as descripted for *S. cerevisiae* strains using minimal media at 30 °C with either ammonium sulfate, arginine, proline, or glutamate as sole nitrogen source.

### Unlabeled and N^15^ labeled metabolomics of *S. pombe* and human plasma

Double Sp-MCUR1 and Sp-CCDC90B and wild-type *S. Pombe* strains were grown for ∼24 h till mid-log phase in minimal media at 30 °C, the pellet was washed two times with PBS and resuspended in Yeast Nitrogen Base (YNB) media without amino acids and ammonium with either ^15^N labeled or unlabeled ammonium sulfate. After 12-16 hours of growth cells were collected, washed three times with water and the pellet was snap frozen in liquid nitrogen and stored at -80°C until use. Yeast cell pellets were resuspended in an adjusted volume of ice-cold 80:20 HPLC grade methanol/water with 100 μl of solvent per 7 mg of cells and subsequently transferred into a 2.0-ml impact-resistant tube containing 300 mg of 1-mm zirconium beads. Samples underwent three 15-s homogenization cycles at 6400 Hz in a Precellys 24® tissue homogenizer. To ensure complete cell lysis, samples were thereafter sonicated for 2 min and vortexed for 30 s. To enable complete protein precipitation, samples were then placed in a −20 °C freezer for 30 min. Samples were thereafter vortexed again for 30 s and centrifuged at 14,000 × g for 10 min at 4 °C. Supernatants were transferred to LC-MS vials containing 200-μl glass inserts. An injection volume of 2.0 μl was used so that ∼140 μg of yeast cells were injected for all samples.

Serum samples were prepared for metabolomics analysis by adding 160 μl of ice-cold HPLC grade methanol to 40 μl of serum. Proteins were precipitated in a -20°C freezer for 30 min followed by vortexing sample for 30 s and centrifugation at 14,000 × g for 10 min at 4 °C. Supernatants were transferred to new vials and dried in vacuo using a Thermo Savant vacuum concentrator at 35 °C. Samples were resuspended in 50 μl of 80:20 HPLC grade methanol/water and transferred to LC-MS vials containing 200 μl glass inserts. An injection volume of 2.0 ul was used for LC-MS analysis.

Yeast cell pellets and serum samples were injected on a Thermo QExactive orbitrap mass spectrometer coupled to a Thermo Vanquish UPLC system. Chromatographic separation was achieved using a Millipore (Sequant) Zic-pHILIC 2.1 × 150mm 5um column maintained at 25 °C with a flow rate of 0.3 mL/min as previously described^24^.

### Wound-healing assays

All cell lines were seeded in 12-well plates and cultured in DMEM with 10% FBS and 2µM uridine until 70% confluency. Then a scratch was made in each well using a 200µl sterile pipette tip of about 50 µm. Cells were washed twice with PBS, to remove the debris and smooth the edge of the scratch and incubated in MEM with 10% FBS and 2µM uridine. Images were acquired every 12h up to 24h after scratch generation using the EVOS XL core (AMEX 1000R, life technologies). Scratch reduction was determined by counting of the cells that cross into the scratch area from their reference point at time 0 and normalized on the number of cells at both the sides of the scratch by Image Pro-Plus 10 software (Media Cybernetics).

### Statistics and reproducibility

Data are represented as mean ± SEM and the statistical analysis of each experiment is described in the figure legends including the statistical tests used and the exact value of biological replicates. For each biological replicate experiment at least 3 technical replicates were used for quantification and data analysis. Normal distribution was tested by Shapiro-Wilk normality test. Statistical analyses were performed using GraphPad Prism (GraphPad Software, version 7).

## Acknowledgements

We acknowledge Dr. Cecilia Garcia-Perez, Dr. Alexandros A Pittis, and Dr. Fabian Hosp for experimental and computational support. FP group was supported by the Munich Center for Systems Neurology (SyNergy EXC 2145 /ID 390857198) and the Initiative and Network Fund of the Helmholtz Association. Work in DM group is supported by Deutsche Forschungsgemeinschaft (grants MO1944/1-2 and MO1944/2-1). DM acknowledges the expert technical assistance of Petra Robisch. TBH and HP received funding from the European Commission (Recon4IMD - GAP-101080997).

## Author contributions

Conceptualization, F.P. and F.V.F.; Methodology, J.V.F., S.B., M.M., K.L., T.H., M.D., H.P.; Formal Analysis, J.W., F.V.F., S.W., Y.C., R.N.; Visualization, J.V.F., and J. W.; Investigation, J.V.F., J.W., V.G, M.S.F., H.C.D., A.L., K.L.; Resources, F.P., M.M., B.S., D.M.; Writing J.V.F, F.P. and F.V.F.; Supervision, F.P., M.J., B.S., D.M., M.M.; Funding acquisition: F.P.

## SUPPLEMENTARY FIGURE LEGENDS

**Figure S1. Validation of MCUR1-StrepII-HA-His_6_.**

**(A)** MCUR1-StrepII-HA-His_6_ is enriched in mitochondria. Analysis of whole-cell (WCL), nuclear (N), cytosolic (C), and mitochondrial (M) fractions isolated from Flp-In^TM^ 293 T-Rex cells after tetracycline induction. Fractions extracts were separated by SDS-PAGE and analyzed by Western blotting. ATP5A, Lamin A/C and β-actin were used as positive controls for mitochondrial, nuclear and cytosolic proteins, respectively.

**(B)** MCUR1-StrepII-HA-His_6_ is an integral mitochondrial membrane protein. Alkaline carbonate extraction at pH 10 and pH 11 of mitochondria isolated from tetracycline-induced Flp-In^TM^ 293 T-Rex cells. Mitochondrial soluble (S) and membrane pellet (P) protein fractions were separated by SDS-PAGE and analyzed by Western blotting. ATP5A and VDAC were used as positive controls for membrane associated and membrane spanning proteins, respectively.

**(C)** Protein topology of MCUR1**-**StrepII-HA-His_6._ Proteinase K (PK) treatment of mitochondria isolated from tetracycline-induced Flp-In^TM^ 293 T-Rex cells. TOM20, TIM23, and cyclophilin F (Cyc F) were used as controls for integral mitochondrial outer membrane, inner membrane and soluble matrix targeted proteins, respectively. T, triton (1%); Dig., digitonin.

In all blots, MCUR1 was detected with anti-HA and anti-MCUR1 antibodies. *Corresponds to untagged MCUR1. IB, immunoblot.

**Figure S2. Knockdown of MCUR1 in HeLa cells.**

MCUR1 protein levels in whole-cell lysates from HeLa cells stably expressing a mitochondria-targeted aequorin (mt-AEQ) and either an empty lentiviral vector (pLKO) or a MCUR1-targeted short hairpin RNA (shRNA; sh1-5). ß-actin was used as loading control.

**Figure S3. MCUR1 deficiency does not impair mt-Ca^2+^ uptake in human cells.**

**(A)** Average traces and quantification of mt-Ca^2+^ transients in controls (C1, C2) and patient (P) skin fibroblasts stably expressing mt-AEQ upon histamine (His) stimulation.

**(B)** Average traces and quantification of mt-Ca^2+^ transients in response to histamine (His) stimulation in control (pLKO) and shMCUR1 HeLa mt-AEQ cells (*n* > 5 biologically independent experiments).

**(C)** Average traces and quantification of mt-Ca^2+^ uptake in digitonin-permeabilized controls and patient skin fibroblasts in presence of glutamate (10 mM) and malate (2 mM) as respiratory substrates.

**(D)** Average traces and quantification of mt-Ca^2+^ uptake in digitonin-permeabilized controls and patient skin fibroblasts in presence of TMPD (0.5 mM)/ ascorbate (2 mM) as respiratory substrates.

**(E)** Average traces of mt-Ca^2+^ buffering capacity in digitonin-permeabilized controls and patient skin fibroblasts in presence of glutamate (10 mM) and malate (2 mM) as respiratory substrates. Ru360 (10 µM) was used as positive control for MCU inhibition (***p<0.001; one-way ANOVA with Tukey’s multiple comparisons test).

**(F)** Average traces of mt-Ca^2+^ buffering capacity in digitonin-permeabilized controls and patient skin fibroblasts in presence of glutamate (10 mM) and malate (2 mM) as respiratory substrates, upon multiple Ca^2+^ spikes.

**Figure S4. MCUR1 deficiency does not alter MCUC stability and assembly in human cells.**

**(A)** Steady-state protein levels of MCUC components in whole-cell lysates from controls and patient skin fibroblasts. ß-actin was used as loading control.

**(B)** Steady-state protein levels of MCUC components in whole-cell lysates from control (pLKO) and MCUR1_sh1 HeLa cells. ß-actin was used as loading control (*n*=3 biologically independent experiments).

**(C)** BN-PAGE analysis of MCUC assembly in mitochondria isolated from controls and patient skin fibroblasts.

**(D)** BN-PAGE analysis of MCUC in mitochondria isolated from control (pLKO) and MCUR1_sh1 HeLa cells. ATP5A was used as loading control (*n*=3 biologically independent experiments). IB, immunoblot.

**Figure S5. Mitochondrial bioenergetics in patient fibroblasts and HeLa MCUR1 knockdown cells.**

**(A)** Measurements of mitochondrial respiration in intact patient (P) and control fibroblasts (C1, C2). (*Left*) Average oxygen consumption rate (OCR) kinetics and (*right*) quantification of basal (State II), non-ADP-stimulated (State IV_o_) and CCCP-stimulated (State III_u_) respiration in response to oligomycin A (O, 1.5 μM) and CCCP (C, 1 μM). Rotenone/antimycin A (R, 2 uM; AA, 4 μM) are used to block RCCI and RCCIII-driven respiration. Data were normalized by total DNA content (*n*=3 biologically independent experiments with more than 4 technical replicates for each condition).

Measurements of RCCI-dependent (**B**) and RCCII-dependent (**C**) OCR in digitonin-permeabilized patient and control fibroblasts. (*Left*) Average OCR kinetics and (*right*) quantification of basal (State II), ADP-stimulated (State III), non-ADP-stimulated (State IV_o_) and CCCP-stimulated (State III_u_) respiration in response to ADP (4 mM), oligomycin A (O, 1.5 μM), and CCCP (C, 15 μM) and in presence of (B) a-ketoglutarate (10 mM)/ malate (10 mM) or (C) succinate (10 mM)/rotenone (2 μM) as respiratory substrates. Rotenone (2 μM) and antimycin A (4 μM) are used as RCCI and RCCIII inhibitors, respectively. Data were normalized by total DNA content (*n*=3 biologically independent experiments, with more than 4 technical replicates for each condition).

**(D)** Measurements of RCCIV-dependent OCR in digitonin-permeabilized patient and control fibroblasts. (*Left*) Average OCR kinetics and (*right*) quantification of basal (State II), ADP-stimulated (State III), non-ADP-stimulated (State IV_o_) and CCCP-stimulated (State III_u_) respiration in response to ADP (4 mM), oligomycin A (O, 1.5 μM), and CCCP (C, 15 μM) and in presence of TMPD (0.5 mM)/ ascorbate (2 mM) as respiratory substrates and antimycin A (2 μM) as RCCIII inhibitor. Azide (Az, 20 uM) was used to block RCCIV-driven respiration. Data were normalized by total DNA content (*n*=3 biologically independent experiments with more than 4 technical replicates for each condition).

**(E)** Measurements of mitochondrial respiration in intact HeLa cells stably expressing a shRNA against MCUR1 (MCUR1_sh1). (*Left*) Average oxygen consumption rate (OCR) kinetics and (*right*) quantification of basal (State II), non-ADP-stimulated (State IV_o_) and CCCP-stimulated (State III_u_) respiration in response to oligomycin A (O, 1.5 μM) and CCCP (C, 1 μM). Rotenone/antimycin A (R, 2 uM; AA, 4 μM) are used to block RCCI and RCCIII-driven respiration. Data were normalized by total DNA content (*n*=3 biologically independent experiments, with more than 4 technical replicates).

**Figure S6. Loss of MCUR1 does not affect the level or assembly of respiratory chain complexes.**

**(A)** Steady-state protein levels of individual RCC subunits in whole-cell lysates from control and patient fibroblasts. ß-actin was used as loading control (*n*=3 biologically independent experiments).

**(B)** Steady-state protein levels of individual RCC subunits in whole-cell lysates from control (pLKO) and MCUR1_sh1 HeLa cells. ß-actin was used as loading control (*n*=3 biologically independent experiments).

**(C)** BN-PAGE analysis RCC assembly in mitochondria isolated from controls and patient fibroblasts. ATP5A was used as loading control (*n*=3 biologically independent experiments).

**(D)** BN-PAGE analysis RCCs assembly in mitochondria isolated from control (pLKO) and MCUR1_sh1 HeLa cells. ATP5A was used as loading control (*n*=3 biologically independent experiments). IB, immunoblot.

**Figure S7. MCUR1 is not required for respiratory growth in *S. cerevisiae*.**

**(A)** Deletion of *FMP32* (*fmp32*Δ) does not result in growth defect on agar plates supplemented with either glucose (YPD) or lactic acid (YPL) and glycerol (YPG) as fermentable and non-fermentable substrates, respectively, when incubated at 24°C, 30°C or 37°C. *FMP32* was deleted by homologous recombination with two different selection cassettes (KAN, kanamycin; HYG, hygromycin) in two different yeast strains (YPH499 and BY4742). Data are representative of two independent experiments.

**(B)** Growth analysis of *WT* and *fmp32*Δ strains in liquid YPD or YPL media at 30°C (mean ± SEM, *n*=4 biologically independent experiments).

**Figure S8. Evolutionary profile of MCUR1 and CCDC90B.**

Phylogenetic distribution of MCUR1 and CCDC90B orthologs across 247 eukaryotes. The phylogenetic tree was reconstructed using the phylogenetic tree generator (https://phylot.biobyte.de/) and visualized using iTOL (https://itol.embl.de/). Homologs of human MCUR1 and CCDC90B were retrieved from ProtPhylo using OrthoMCL with more than 0% match length and inflation index of 1.1 for orthology assignment.

**Figure S9. Effect of MCUR1 knockout in yeast and HeLa cells.**

Growth of prototrophic wild-type (*WT*) and *Sp-MCUR1* knockout *S. pombe* cells in minimal media with either arginine (**A**) or glutamate (**B**) as sole nitrogen sources.

**(C)** Rescue of *Sp-MCUR1* knockout growth defect in minimal media with ammonium sulfate by supplementation with glutamate.

**(D)** Steady-state MCUR1 levels in whole-cell lysates from controls and patient fibroblasts compared with patient fibroblasts expressing full-length MCUR1 (Rescue). ß-actin was used as loading control.

**(E)** Proteomics profiling in shMCUR1 Hela cells. The Volcano plot shows significantly different proteins (black filled circles) between control and MCUR1 deficient cells with *p* values below 0.05. CCDC90B was identified only in control cells (FDR 0.05, S0 0.1).

## List of Abbreviations

AEQ: Aequorin
BN–PAGE: Blue native–polyacrylamide gel electrophoresis
Ca^2+^: Calcium ion
CCCP: Carbonyl cyanide m-chlorophenylhydrazone
CCDC: Coiled-coil domain-containing
CCDC90A: Coiled-coil domain-containing protein 90A, also known as MCUR1
CCDC90B: Coiled-coil domain-containing protein 90B
COX: Cytochrome c oxidase
DNA: Deoxyribonucleic acid
EMT: Epithelial–mesenchymal transition
ER: Endoplasmic reticulum
EGTA: Ethylene glycol-bis(ß-aminoethyl ether)-N,N,N’,N’-tetraacetic acid
FARS2: Phenylalanyl-tRNA synthetase 2, mitochondrial
FMP32: Fission Mitochondrial Protein 32
gnomAD: Genome Aggregation Database
HA: Hemagglutinin
HEK293: Human embryonic kidney 293
IARS2: Isoleucyl-tRNA synthetase 2, mitochondrial
KO: Knockout
LC–MS/MS: Liquid chromatography–tandem mass spectrometry
MCU: Mitochondrial calcium uniporter
MCUC: Mitochondrial calcium uniporter complex
MCUR1: Mitochondrial calcium uniporter regulator 1
MRI: Magnetic resonance imaging
mt-Ca^2+^: Mitochondrial calcium
mRNA: Messenger ribonucleic acid
OCR: Oxygen consumption rate
OXPHOS: Oxidative phosphorylation system
RCC: Respiratory chain complex
RCCI: Respiratory chain complex I
RCCIV: Respiratory chain complex IV
RNA: Ribonucleic acid
SDH: Succinate dehydrogenase
SERCA: Sarco/endoplasmic reticulum Ca²?-ATPase
shRNA: Small hairpin RNA
Sp-MCUR1: *Schizosaccharomyces pombe* mitochondrial calcium uniporter regulator 1
Sp-CCDC90B: *Schizosaccharomyces pombe* coiled-coil domain-containing protein 90B
TAP: Tandem affinity purification
TCA: Tricarboxylic acid
TORC1: Target of rapamycin complex 1
VARS2: Valyl-tRNA synthetase 2, mitochondrial
WARS2: Tryptophanyl-tRNA synthetase 2, mitochondrial
WT: Wild-type

