## Supplementary figures and images for "MCUR1–CCDC90B complex is a conserved mitochondrial scaffold regulating metabolic homeostasis"

### Supplementary figures

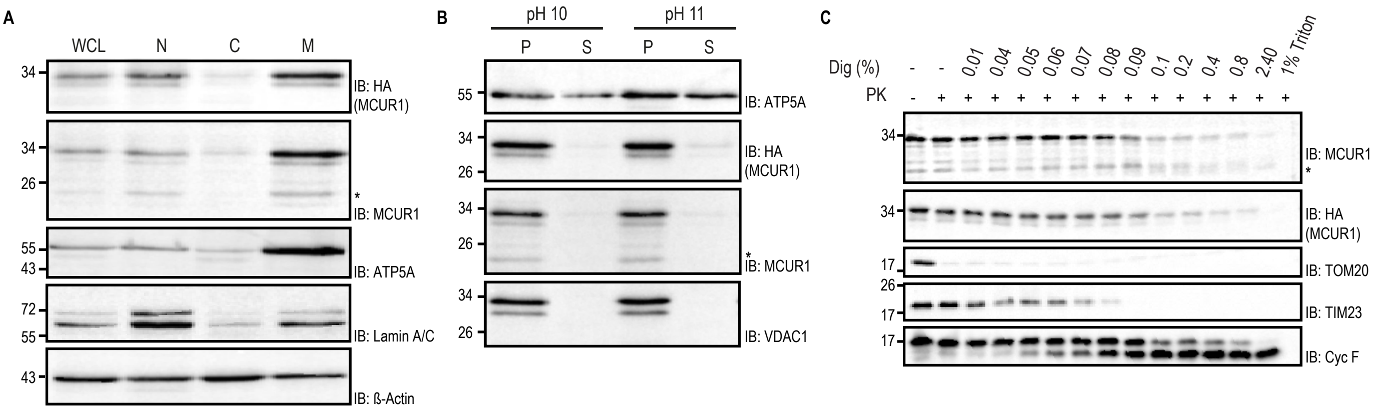

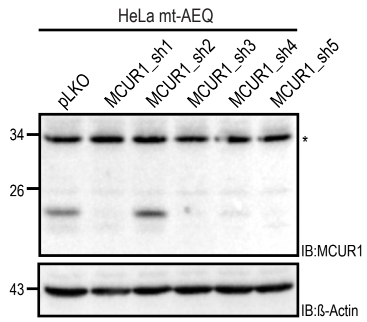

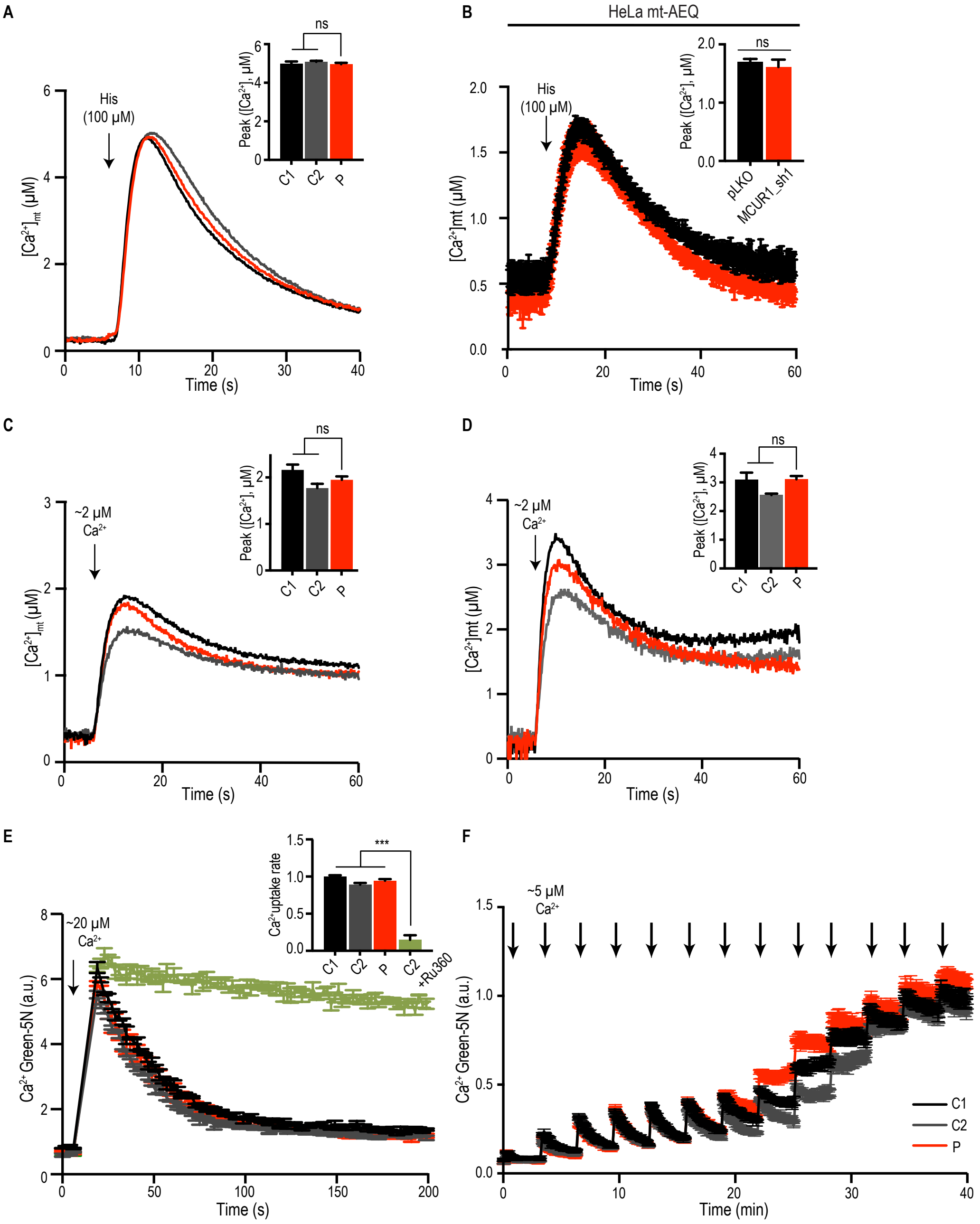

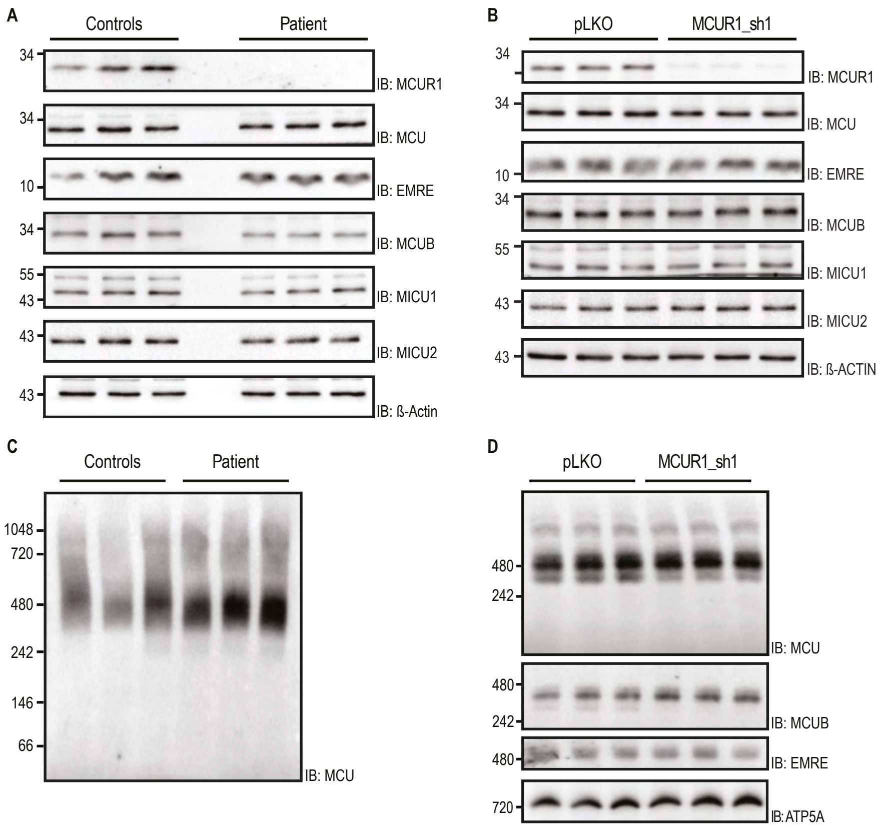

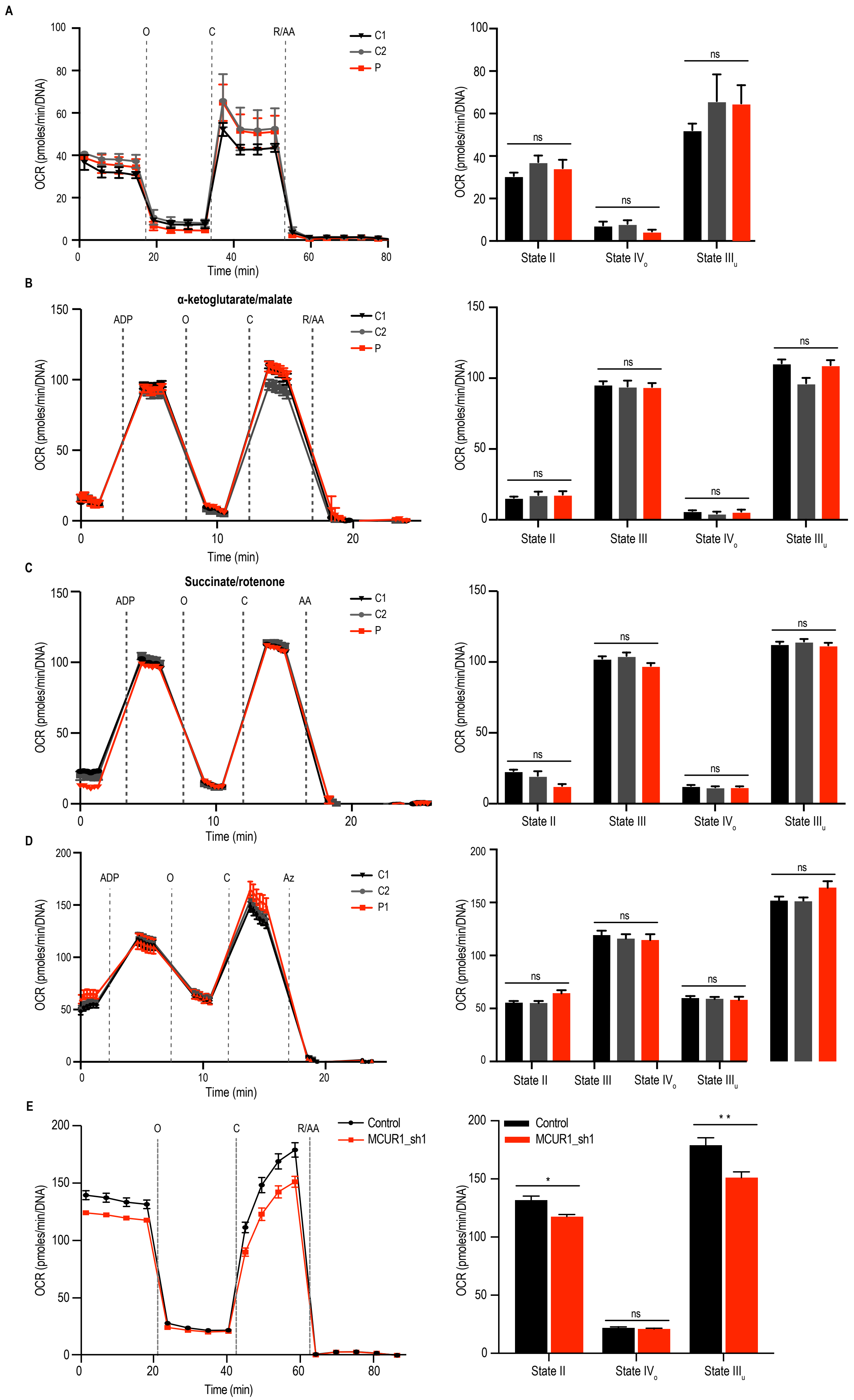

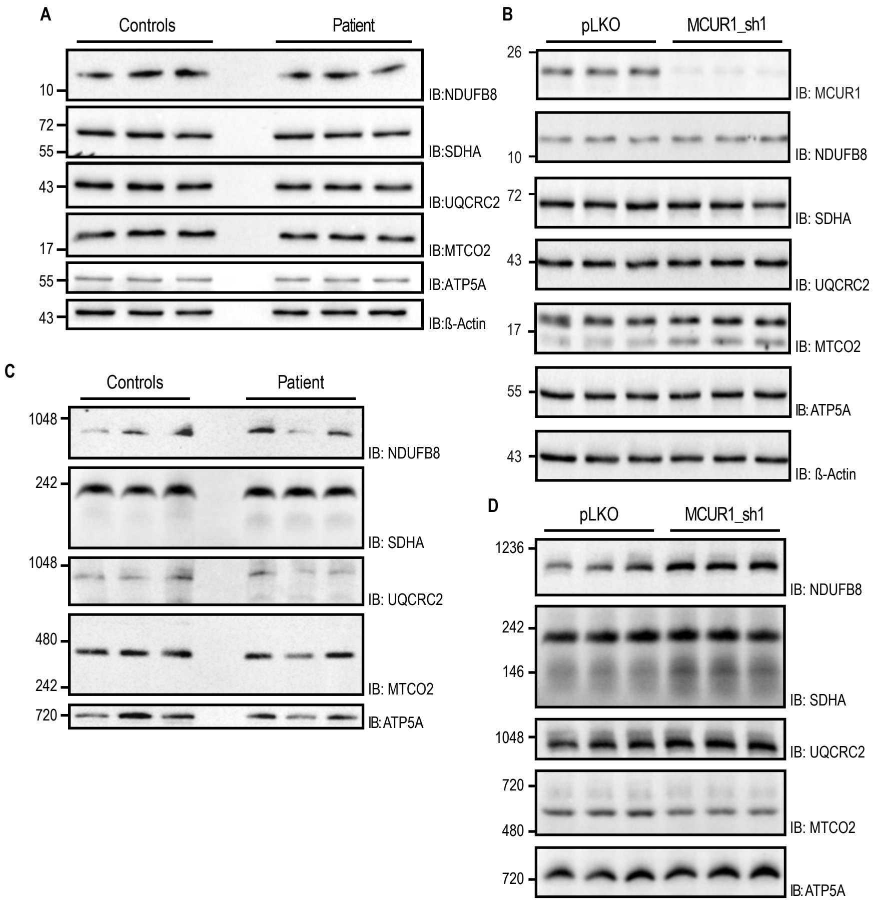

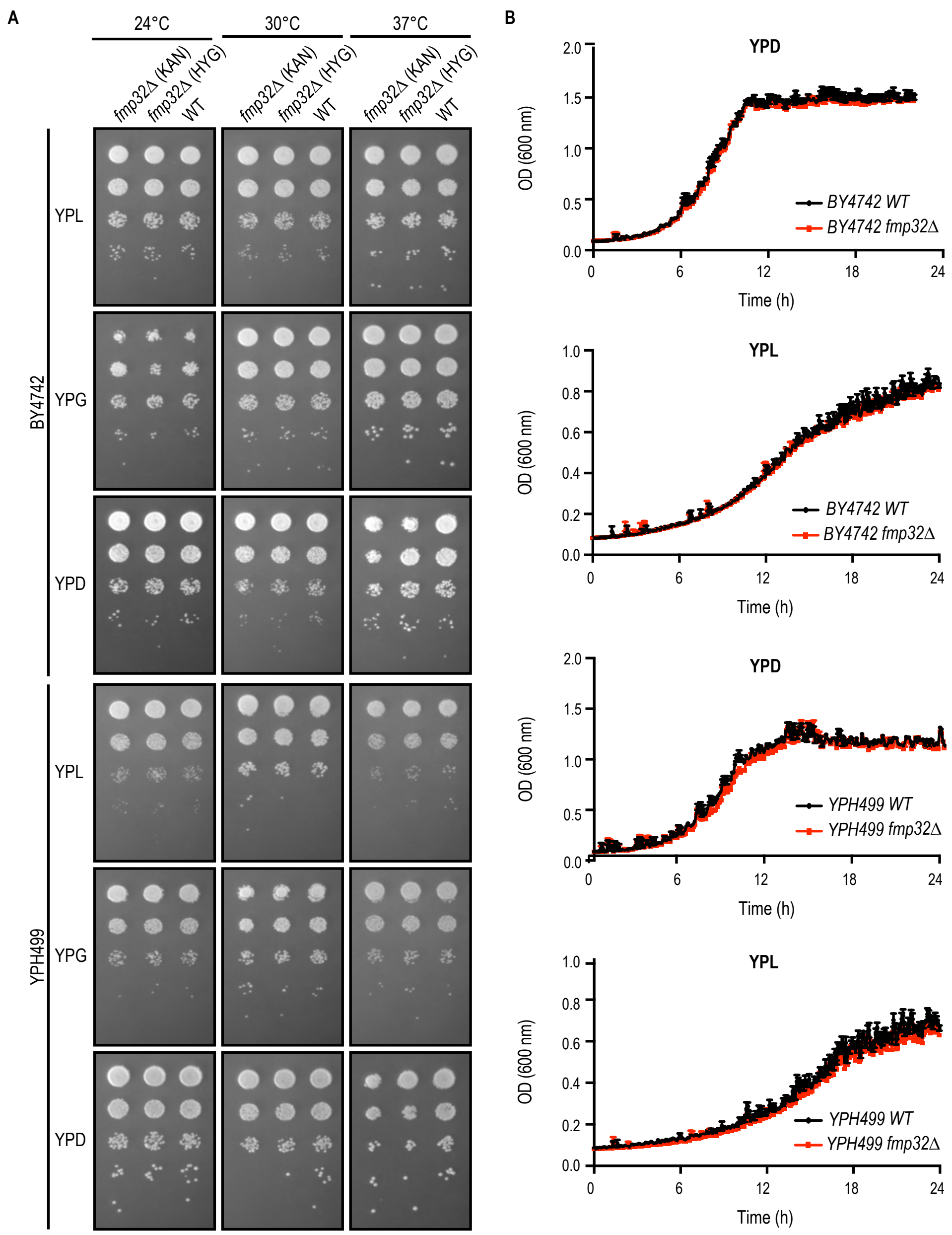

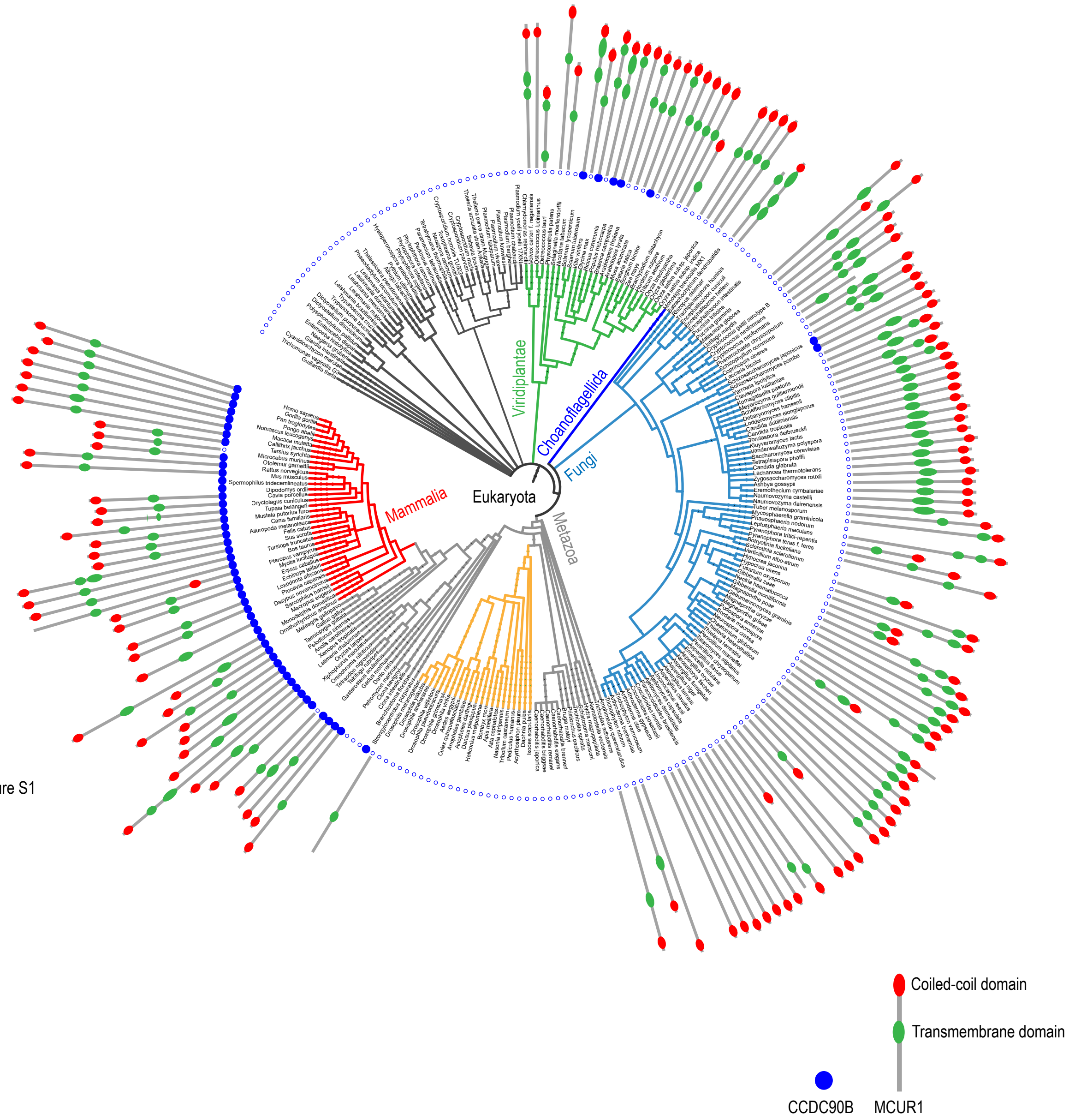

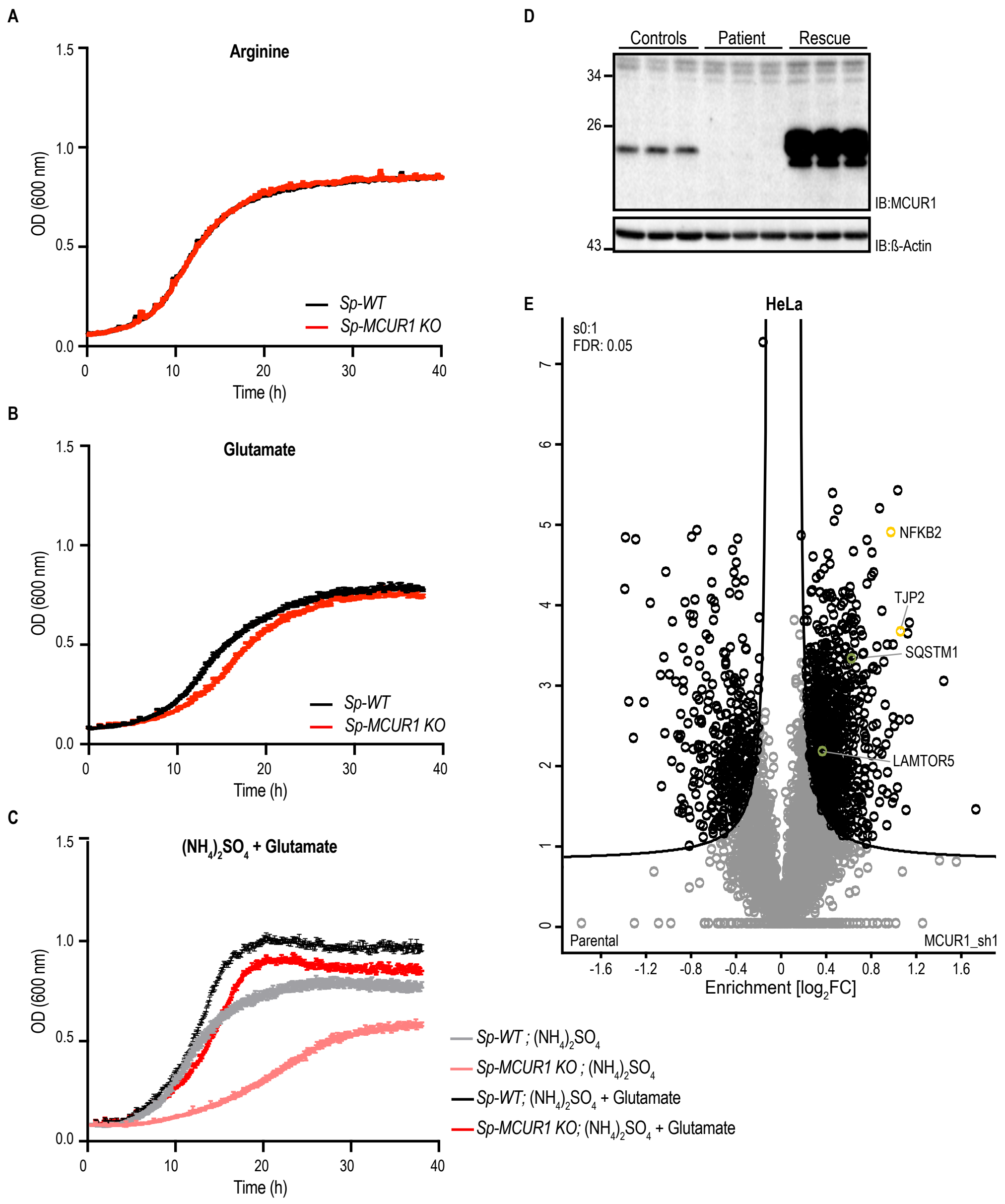
